# Enhancer Dynamics and Spatial Organization Drive Anatomically Restricted Cellular States in the Human Spinal Cord

**DOI:** 10.1101/2025.01.10.632483

**Authors:** Elena K. Kandror, Anqi Wang, Mathieu Carriere, Alexis Peterson, Will Liao, Andreas Tjärnberg, Jun Hou Fung, Krishnaa T. Mahbubani, Jackson Loper, William Pangburn, Yuchen Xu, Kourosh Saeb-Parsy, Raul Rabadan, Tom Maniatis, Abbas H. Rizvi

## Abstract

Here, we report the spatial organization of RNA transcription and associated enhancer dynamics in the human spinal cord at single-cell and single-molecule resolution. We expand traditional multiomic measurements to reveal epigenetically poised and bivalent active transcriptional enhancer states that define cell type specification. Simultaneous detection of chromatin accessibility and histone modifications in spinal cord nuclei reveals previously unobserved cell-type specific cryptic enhancer activity, in which transcriptional activation is uncoupled from chromatin accessibility. Such cryptic enhancers define both stable cell type identity and transitions between cells undergoing differentiation. We also define glial cell gene regulatory networks that reorganize along the rostrocaudal axis, revealing anatomical differences in gene regulation. Finally, we identify the spatial organization of cells into distinct cellular organizations and address the functional significance of this observation in the context of paracrine signaling. We conclude that cellular diversity is best captured through the lens of enhancer state and intercellular interactions that drive transitions in cellular state. This study provides fundamental insights into the cellular organization of the healthy human spinal cord.

## INTRODUCTION

The human spinal cord is the principal conduit for somatosensory input and motor output, enabling voluntary and autonomic movements. To support these functions, neurons and glia are patterned across two anatomic axes: rostrocaudal and dorsoventral. The rostrocaudal axes are defined by vertebral segments, along which motor neurons are arranged in columns to support control of the arm (cervical), axial (thoracic), and leg (lumbar) muscles. The dorsoventral axis is characterized by Rexed laminae, in which stereotyped neuronal subtype cytoarchitecture controls discrete sensorimotor processing steps. While the patterning of neurons across the spinal cord is well established, the question of how glial cells respond to the local demands of neural circuitry remains unclear. The cellular organization of the spinal cord is similar between the thoracic and lumbar regions, yet differences in cellular responses emerge in ALS^1^ and cancer^2^. These differences may result from anatomic differences in glial reactivity to pathological states. We reasoned that such responses arise from cell type-specific differences in gene regulation prompted by intercellular signaling. Combinatorial patterns of gene regulation may establish distinct glial cellular states along the rostrocaudal axis of the spinal cord, explaining how motor circuits may be differentially supported. Furthermore, cellular subtypes may have the potential to access different physiological states, depending on their anatomical position in the spinal cord and differences in specific communal cellular interactions. Regulatory plasticity, defined as a cell’s ability to transition to an altered cellular state, arises from the convergence of chromatin state, encoded genetic determinants, in concert with induction by autocrine and paracrine signaling^3^. Transcriptional activation, therefore, is a consequence of cellular induction, followed by orchestrated changes in chromatin valence, transcription factor binding to cis-regulatory DNA elements, and long-range enhancer interactions with promoters. The capacity to orchestrate such transitions constitutes an additional dimension of cellular diversity, driven by poised enhancer states and complex cell-cell interactions. Cellular diversity can thus be recast in the context of cellular plasticity and locally interacting networks of cells that provide environmental cues to trigger cellular state changes.

Here, we characterize the transcriptional, epigenetic, and spatial diversity of neurons and glia in the human spinal cord, define the regulatory logic that enables their specification, and uncover a fundamental epigenetic mechanism for gene activation that enables specialized function. We consider transcriptional activation in the context of dynamic changes in chromatin states, transcription factor binding to distal regulatory elements, and engagement with the basal transcriptional machinery^4^. Additional regulatory mechanisms include dynamic patterns of DNA methylation and transcription factor activity^5^. Remarkably, poised enhancer states and distal regulatory elements with unrealized transcriptional potential are retained following development, yielding differences in cellular plasticity^6^. Initially, we identified enhancer dynamics at the single cell level, revealing regulatory strategies and differential gene expression patterns associated with cellular identity and anatomically defined constraints in cellular states between the thoracic and lumbar spinal segments. We then defined cellular subtypes based on common regulatory variation and pinpointed previously undetected active enhancers in the absence of chromatin remodeling with cell type and anatomic specificity. These observations provide novel insights into the cellular organization of the human spinal cord in the context of segment-level enhancer dynamics.

Finally, we introduce the detection of cellular networks as a third level of spatial organization: repeat patterns of cell types that recurrently exist in proximity to one another, tile throughout a cross-section of the spinal cord, and are likely responsible for self-contained paracrine signal transduction in the central nervous system. We developed an approach to project high-depth transcriptomic measurements onto single cell and spatially resolved multiplexed *in situ* profiled spinal cord sections. These spatial data were used to identify neighborhoods of interacting cells. Analysis of complementary receptor-ligand pairs shared by cells within a neighborhood made it possible to link molecular induction with the regulatory capacity of these populations. Ultimately, these mechanisms lead to the formation of interacting cellular communities that support physiological function with anatomic specificity. Taken together, our findings recast cellular identity within the human spinal cord through the lens of regulatory plasticity and anatomic organization. Our analyses provide a molecular and cellular template by which future studies of neurodegenerative diseases can be compared.

## RESULTS

### Cellular Diversity is Driven by Restricted Regulatory Logic in the Human Spinal Cord

The stoichiometry of neurons and glia in the healthy human spinal cord was previously described^7,8^, but the regulatory logic underlying cellular specification, and importantly, the programs that are requisite for transitions to altered states during disease, have not been assessed. We, therefore, started by characterizing cellular heterogeneity and the underlying transcription factor activity differences between cellular subtypes in the spinal cord. We profiled gene expression and chromatin accessibility in 150,000 nuclei from the thoracic (T4) and lumbar (L4) regions of six healthy donor spinal cords (Figure 1A). Nuclei from Donor 1 were processed independently for gene expression and chromatin accessibility, while nuclei from Donors 2-6 were profiled through simultaneous multiomic measurements. We detected an average of 2000 genes and 7000 fragments per nuclei (Supplementary Figure 1A). To minimize the deleterious effects of a postmortem interval, we established a surgical procedure in which spinal cord tissue was obtained from organ donors within 60 minutes of tissue donation (Donors 2-6), thereby minimizing artifactual transcriptional changes observed in hypoxic conditions. We integrated multiomic data across spinal cord segments and identified 37 major neuronal and glial cell populations resident in the healthy spinal cord. The postmortem interval effect from Donor 1 resulted in a sharp decrease in neuronal recovery but had a negligible impact on cluster composition. The observed glial and neuronal cell type classifications are consistent with the expected stoichiometry of cell types in the adult human spinal cord and between donors and segment levels (Supplementary Figure 1B,C).

**Fig. 1.**
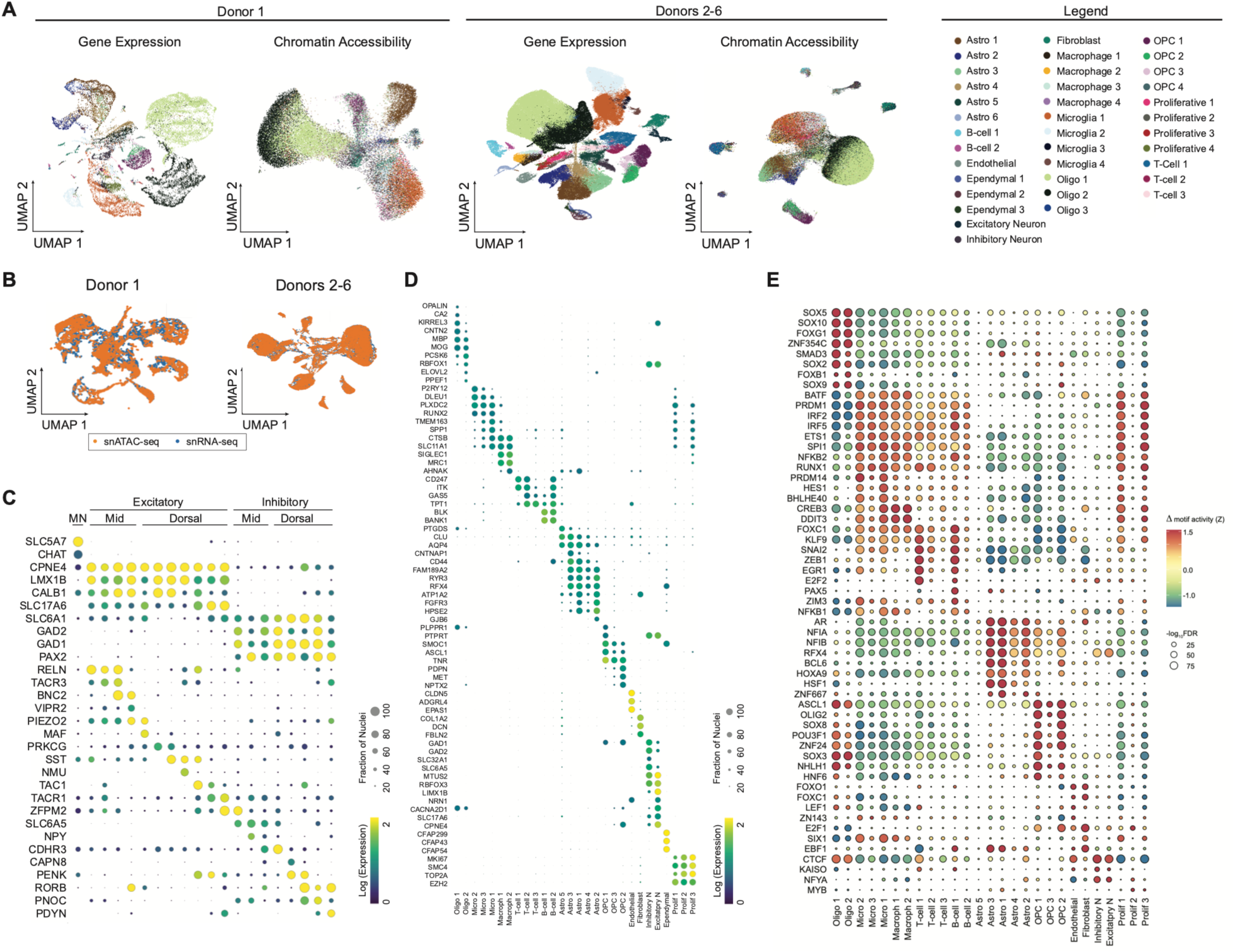
Regulatory Logic and Transcriptional Activity in the Adult Human Spinal Cord. **(A)** UMAP embeddings of independently measured snRNA and ATAC from (left) 40,000 nuclei from T4 and L4 spinal segments of a postmortem donor and (right) 100,000 simultaneously profiled nuclei from T4 and L4 spinal segments of five organ donors. Stoichiometry of cell types is consistent between donors and modalities. **(B)** Optimal transport-based integration of snRNA-seq (blue) and snATAC-seq (orange) yields accurate co-embedding of data into a shared multimodal feature space. **(C)** Subclustering of neurons based on gene expression reveals expected cholinergic, excitatory, and inhibitory populations spanning the dorsoventral axis, as shown by marker gene expression in the dot plot. **(D)** Unbiased clustering of single nuclei gene expression reveals heterogeneous cellular subtypes, which can be identified by marker gene expression as shown in the dot plot. **(E)** Motif activity analysis of aggregated snATAC measurements reveals discrete regulatory programs in cellular subtype specification.

Cross-modal data integration requires that datasets be merged into a shared feature space, which can be challenging without a priori known anchor genes. We therefore developed an unsupervised computational strategy based on optimal transport^9^ to harmonize transcriptional readout with chromatin accessibility. To establish this approach, we separated simultaneous assays of chromatin accessibility and transcriptome derived from individual nuclei, in order to generate a ground truth synthetic dataset. Multiomic data from each case were aggregated and subjected to canonical correlation analysis and singular value decomposition for coarse co-embedding into a shared feature space. We then utilized entropically regularized optimal transport, minimizing the Wasserstein distance associated with pairing chromatin accessibility with snRNA-seq data. This approach accurately co-embedded data, surpassing the accuracy observed in existing methods (Supplementary Figure 1D). Nuclei from Donors 2-6, which had been simultaneously profiled for RNA and chromatin accessibility, show near total integration (Figure 1B). Donor 1, in which separate nuclei preparations were independently measured at higher depth, shows similarly high concordance between modalities. For downstream analysis, we focused on nuclei from Donors 2-6 (deceased transplant organ donor tissue). We subclustered the neurons and identified 20 populations of cholinergic, excitatory, and inhibitory neurons spanning the dorso-ventral axis of the spinal cord, consistent with the known organization of neurons in the Rexed laminae (Figure 1C).

We observed extensive heterogeneity in the basal transcriptional state of glial cells in the healthy spinal cord (Figure 1D, Supplementary Table 1). Oligodendrocytes are divided into two dominant populations, Oligo1 (OPALIN, CA2) and Oligo2 (KLK6, ELOVL2). Histologically, oligodendrocytes can be defined by their preferred myelination targets: multiple thin axons or a dedicated large axon^10^. We speculate that the enriched myelination program in Oligo2 may predispose this population to the support and maintenance of thick, descending tract axons. We also observe a small population of Olig3 (ENPP6), which corresponds to a rare population of newly formed oligodendrocytes in the adult human spinal cord. Oligodendrocyte progenitor cells (OPCs) are evenly distributed across three main clusters: synapse-associated OPC1(PTPRT), migratory OPC2 (MET), and resting OPC3 (TNR). A rare population of OPC4 shares transcriptional signatures with OPC1 and OPC2 and may represent a transition state between the two. Healthy microglia are predominantly distributed across three states: phagocytotic Micro1 (SPP1), scavenging Micro2 (P2RY12), and resting Micro3 (PLXDC2). In addition, we observed two populations of proliferating microglia, Prolif1 corresponding to actively dividing Micro2, and Prolif3 corresponding to actively dividing Micro1, suggesting that the physiological state of a progeny microglial cell is determined by the state of its parent cell. Astrocytes are distributed across 4 major populations: fibrous Astro1 (AQP4), protoplasmic Astro2 (GJB6), paranodal Astro3 (CNTNAP1), and Astro4 (RFX4), along with a rare population of regulatory Astro5 (PTGDS). We also identified populations of fibroblasts (COL1A2), endothelial (CLDN5), and ependymal (CFAP299) cells. Finally, infiltrating B-cells (BLK) and T-cells (ITK) were observed predominantly from a single donor, while a small population of macrophages (MRC1) was distributed across Donors 2-6. The coverage of this dataset spans the cell types known to be resident in the adult spinal cord and identifies the heterogeneous nature of resting glial states.

We then identified the cis regulatory programs that govern cell type specification and maintenance of glial and immune cells by leveraging the stereotyped architecture genes, based on chromatin accessibility 100 kb upstream of transcription start sites as the range for enhancer element detection. For each gene enriched in a cluster, we calculated the motif activity of a comprehensive panel of transcription factors (TFs), which serve as a computational proxy for TF participation in cell type-specific gene regulation (Figure 1E). Microglia, derived from the yolk sac and sharing a common lineage with macrophages^11^, have a distinct regulatory profile from other glial cells capable of producing a characteristic immune response (SPI1, IRF2, PRDM1). Due to their shared precursors, microglia and macrophages share a regulatory logic with macrophages, differing only through the unique activity of BHLHE40, a core regulatory transcription factor required for lipid clearance^12^. T cells and B cells, infiltrating immune cells, are modulated by a distinct immune program, including E2F2 and PAX5. Astrocytes are uniquely enriched for HSF1 activity, a repressor of neurotoxic reactivity^13^. They are also strongly enriched for the canonical regulatory factor X (RFX) and nuclear factor I (NFI) transcription factor motifs, both of which display a significant overlap with OPCs. OPCs can differentiate into both oligodendrocytes and protoplasmic astrocytes^14^, which may explain their shared astrocytic regulatory program *in vivo*. OPCs also share common motif activity with oligodendrocytes driven by the bHLH transcription factor ASCL1 but display a stronger reliance on OLIG2 and SOX8 than their mature counterparts. These two transcription factors are critical for remyelination programs^15,16^, and their preferential activity in OPCs rather than mature oligodendrocytes suggests the importance of adult-generated OPCs in recovery from demyelinating disorders. Both oligodendrocyte subtypes are uniquely regulated by SOX2, SOX9, and SOX10, transcriptional programs responsible for their terminal differentiation and myelination capacity^17-19^. Taken together, these data provide a view of the complex landscape of cell types in the healthy human spinal cord, along with the underlying regulatory logic that maintains their committed cellular states.

### Spinal Enhancer Dynamics Operate in the Absence of Chromatin Potential

Epigenetic regulation of gene expression is only partially governed by histone displacement from regulatory sequences, as evidenced by poor concordance between transcriptional readout and ATAC measurements at proximal regulatory regions^20-23^. We, therefore, sought to identify a complementary regulatory strategy that explains cell type specification. Histone modifications are an evolutionarily conserved mechanism for defining active and silenced chromatin regions and, as such, are drivers for transcriptional activation and silencing. Active enhancers and promoters are marked by Histone H3 acetylation (H3K27ac), while gene repression is marked by Histone H3 trimethylation (H3K27me3). In combination with these two modifications, a third histone mark, H3K4me1, defines bivalent or poised sites that are primed for changes in activation. Here, we show that integrating measurements for these modifications with chromatin accessibility enables the identification of cryptic enhancers that function independently of histone displacement and regulate genes critical for cellular specification.

To study this phenomenon in the human spinal cord, we developed an approach that makes it possible to simultaneously measure ATAC-seq and histone modifications in individual nuclei. This method leverages a single nuclei sequential antibody directed barcoded tagmentation assay (Sequential Tagmentation with Barcoded Sequencing, STAB-seq) directed against histone modifications (H3K27ac, H3K4me1, H3K27me3) which, when analyzed in tandem, identify bivalent active (H3K27ac/H3K4me1), bivalent poised (H3K27me3/H3K4me1), primed (H3K4me1), and silenced (H3K27me3) proximal and distal regulatory elements^6,24-28^. Antibodies specific for these modifications were incubated with spinal cord nuclei, followed by treatment with secondary antibody and incubation with Protein A-Tn5 loaded with calling cards (barcoded transposable elements). We followed antibody-directed tagmentation with a general tagmentation using a transposon complex lacking calling cards (Figure 2A). Thus, the assay simultaneously detects the enhancer state alongside all accessible chromatin, providing cell-type-specific information. We profiled ∼90,000 nuclei isolated from the T4 and L4 spinal cord segments from two deceased transplant organ donors, conducting assays for H3K27ac, H3K4me1, and H3K27me3, followed by a non-barcoded tagmentation. We detect an average of 1600 fragments per nuclei from reads containing calling cards, and 3200 fragments per nuclei for unbarcoded reads (Supplementary Figure 2A). The introduction of histone modification-specific calling cards in STAB-seq does not interfere with unbarcoded ATAC profiles, which are consistent between all three modifications profiled (Supplementary Figure 2B). These data were integrated utilizing optimal transport, with cellular identity robustly detected by unbarcoded chromatin accessibility (Figure 2B, Supplementary Figure 2C). Aggregate tracks identified the presence of bivalent active and poised chromatin with mutual exclusivity (Figure 2C, Supplementary Figure 2D). Regulatory regions for genes specifically expressed in oligodendrocytes, OPCs, microglia, and astrocytes show different acetylation profiles between the cell types, defining cellularly distinct transcriptionally active chromatin (Figure 2D).

**Fig. 2.**
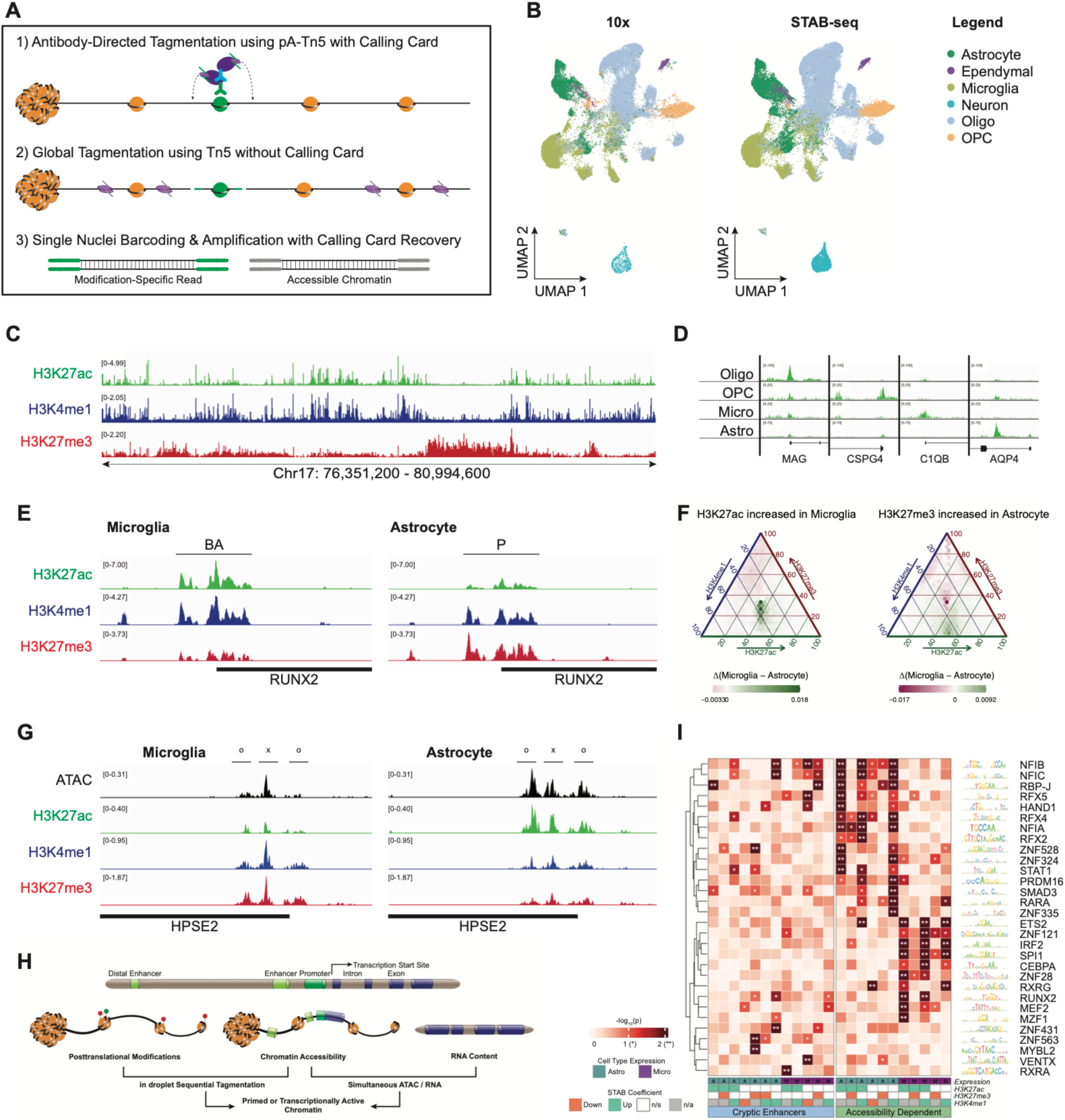
Enhancer Profiling Dissects Chromatin Potential from Accessiblity Independent Regulatory Activity. **(A)** Experimental workflow of Sequential Tagmentation with Barcoded Sequencing (STAB-seq) for the introduction of a histone modification-specific calling card prior to traditional unbarcoded ATAC. **(B)** UMAP representations of (left) 10X-based single nuclei chromatin accessibility and (right) unbarcoded STAB-seq chromatin accessibility show the equivalent distribution of major cell types between modalities. **(C)** Aggregated STAB-seq reads for H3K27ac (green), H3K4me1 (blue), and H3K27me3 (red) along a 4.6Mb track of chromosome 17, demonstrating mutually exclusive active or silent chromatin regions. **(D)** STAB-seq reads for H3K27ac aggregated by cell type show selectively active enhancers at cell type-specific marker genes. **(E)** At the RUNX2 locus, astrocytes and microglia show an inverse relationship between activating and repressing histone modifications: Bivalent Active (BA, H3K27ac+H3K4me1) in microglia and Poised (P, H3K27me3+H3K4me1) in astrocytes. **(F)** Triangle plot demonstrating a consistent elevation of H3K27ac and decrease in H3K27me3 in enhancers that are active in microglia (left) and silenced in astrocytes (right). **(G)** Out of three regulatory peaks at the HPSE2 locus, two show concordant increases in ATAC accessibility and H3K27ac signal in astrocytes vs microglia (o, consistent with chromatin remodeling), while one demonstrates increased H3K27ac signal in astrocytes absent a significant change in ATAC accessibility between cell types (x, consistent with a cryptic enhancer). **(H)** Schematic of 3 modality integration of snSTAB-seq, snRNA-seq, and snATAC-seq, enabling peak-gene associations, regulatory site discovery, and transcription factor analyses. **(I)** Matrix plot showing the significance of TFA scores between astrocytes and microglia organized by TF preference for cryptic vs. remodeling-dependent enhancers between cell types. Peak-associated gene expression was calculated to be enriched in astrocytes (A) or microglia (M). The STAB coefficient, defined as the level of modification enrichment at TF-associated peaks between the two cell types, was calculated as enriched (up), depleted (down), not significant (n/s), or not considered (n/a).

We first asked if histone valence, the contribution of activating versus repressive histone modifications, at a gene regulatory region can define a glial state. We focused on RUNX2, a pioneering transcription factor that has the capacity to alter chromatin accessibility in regulatory regions of its downstream target genes^29^. RUNX2 is constitutively expressed in microglia^30^, inhibiting ameboid transitions^31^. While RUNX2 is normally not expressed in astrocytes, its activation is critical for the suppression of astrocytic reactivity and scarring after injury or immunological challenge^32^. This pattern of expression suggests that RUNX2 should have a positive valence in microglia and incomplete repression (facultative as opposed to constitutive silencing) in astrocytes. The histone code supporting this model of regulation is bivalency: bivalent active (BA) in microglia and bivalent poised (P) in astrocytes. Our STAB-seq results revealed that this is, in fact, the exact mechanism for RUNX2 regulation in glial populations of the spinal cord (Figure 2E). We considered whether this regulatory pattern occurs more generally between astrocytes and microglia and identified combinatorial changes in activating and repressive histone modifications at regulatory sites for hundreds of genes within these glial subtypes (Figure 2F).

We then asked if altered chromatin accessibility is a prerequisite for histone valence to impact gene expression. We highlight HPSE2, which is specifically expressed in astrocytes and prevents the clearance of plaques from the CNS^33^. Given its cell type-specific expression, it would be reasonable for the HPSE2 regulatory elements to be uniquely accessible in astrocytes. Remarkably, while this is true for two proximal regulatory regions for HPSE2 (o), we found that a third region is equally accessible in both glial types (x) (Figure 2G). This peak is uniquely regulated by chromatin valence–acetylation in astrocytes and trimethylation in microglia. We refer to such sites as cryptic enhancers, regulatory regions of the genome that are governed not by chromatin remodeling but rather by histone valence. Cryptic enhancers contribute to the dissonance between the observed expression of a gene and its calculated gene activity score based on chromatin accessibility.

We extended our analysis to determine if distal elements contribute to transcriptional activation by using optimal transport to integrate single nuclei multiomic assays for chromatin accessibility and RNA transcription with our STAB-seq results. ATAC-seq data from both measurements were used by optimal transport to anchor our results. This approach offers a platform to directly connect changes in chromatin accessibility, enhancer state, and gene activation (Figure 2H, Supplementary Figure 2E). We then modified and employed Scarlink^34^, a linear regression model, that makes possible the detection of changes in chromatin modification states, accessibility, and their impact on transcriptional activation. We observed wholesale changes in histone modifications in cell type specific enhancers, revealing changes in bivalency in the absence of chromatin remodeling. Finally, we asked if transcription factor activity (TFA) at distal enhancers was biased towards accessibility dependent versus independent regulation. The results across microglial and astrocyte populations identify transcriptional enhancer classes that impact glial gene activation, which is not detected in traditional multiomic analyses (Figure 2I). The NFIX family of transcription factors (TFs) activate astrocytic genes during development^35^, which include genes that mediate homeostasis. We observed a requirement for chromatin reorganization 100kb upstream of target genes for these TFs. However, these target genes are activated in microglial populations only in enhancers in which histones are constitutively displaced. Conversely, Regulatory Factor X (RFX) TFs, which impact the regulation of a broad range of genes, including MHC class II genes^36^, show significant activity in microglia in the absence of chromatin remodeling but require remodeling in astrocytes. Surprisingly, these are not recognition sites for canonical pioneering transcription factors. Instead, they constitute a novel class of valence-dependent transcription factor binding sites that leverage cryptic enhancers to regulate gene expression. Taken together, these findings demonstrate previously unappreciated enhancer dynamics that operate with cell type specificity.

### Distinct Cell Type Specific Gene Regulatory Networks in the Thoracic and Lumbar Segments

While the neuraxial distribution of motor neuron columns has previously been well described^37-39^, the accompanying glial diversity across spinal segments has not been examined. Statistics from clinical studies in ALS^40-43^ and cancer^2^ suggest that cells between the thoracic and lumbar regions have inherently different responses to disease pathology. However, the underlying molecular basis for this is not understood. We reasoned that such responses are a consequence of differences in regulatory plasticity along the rostrocaudal axis. To understand the inherent differences between glial cell regulatory dynamics between the T4 and L4 segments of the spinal cord, which may obscured during coembedding (Supplementary Figure 3A), we separated the nuclei from each segment and independently performed differential gene expression and accessibility measurements to identify cell types, which remained consistent with the labels from the joint space. We used optimal transport to align the profiles from each modality and identified the nuclei and clusters with shared features (Figure 3A). To minimize uninformative signal from spurious background accessibility changes, we further used optimal transport to project active chromatin signal, identified by H3K27ac peaks, onto the data from segment isolated nuclei.

**Fig. 3.**
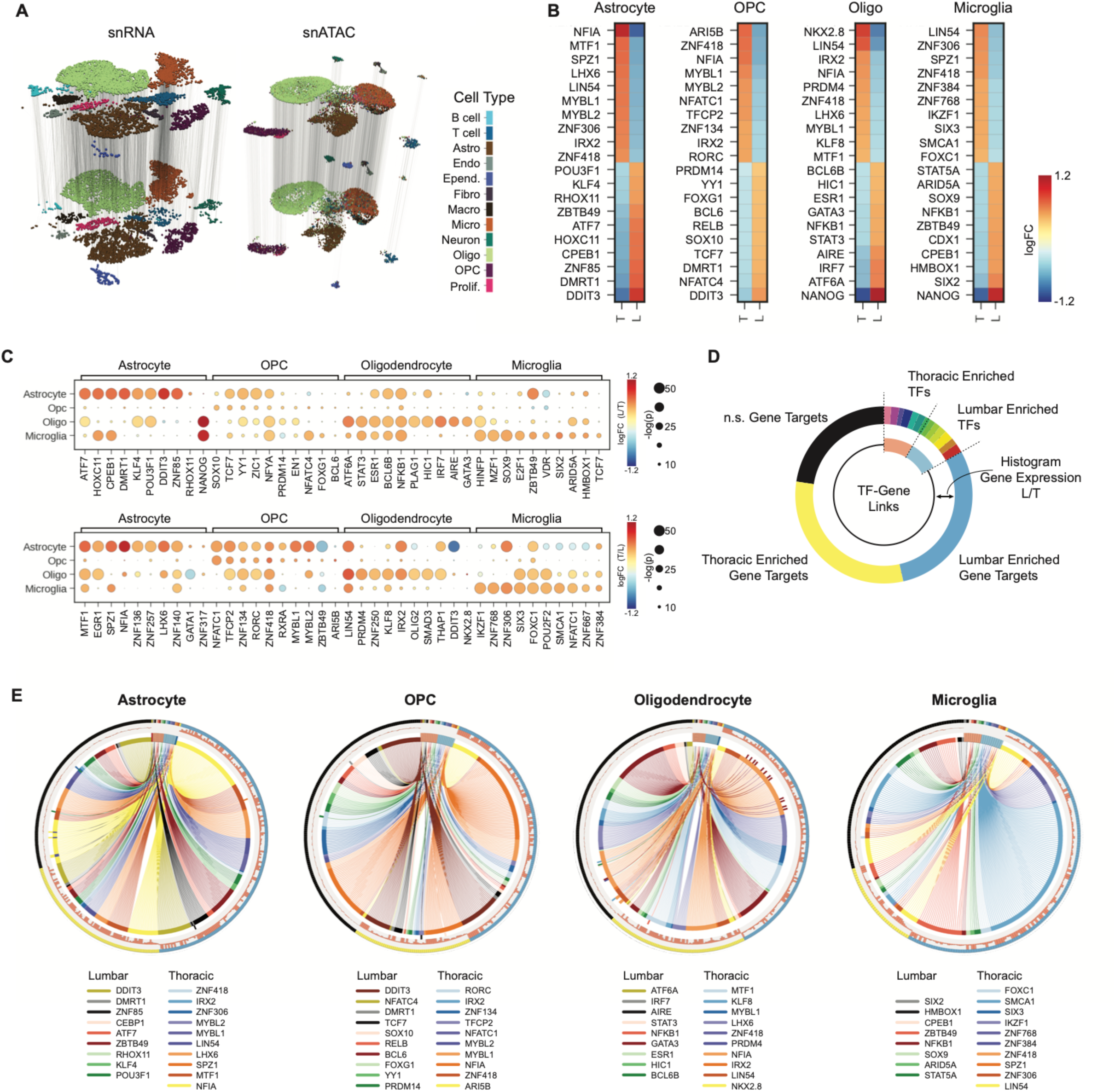
Distinct Regulatory Patterns of Thoracic and Lumbar Glial Cells. **(A)** Nuclei from organ donors profiled simultaneously for snRNA and snATAC were aggregated and clustered separately from the T4 (top) and L4 (bottom) regions of the spinal cord to retain biological variation. Optimal transport was used to align RNA and ATAC measurements between segments. **(B)** Differential motif enrichment in the T4 and L4 regions of the spinal cord across the four major glial populations shows robust differences between segments. **(C)** Dot plot of cell type specific motif activity enrichment in the L4 (top) and T4 (bottom) segments of the adult human spinal cord demonstrate distinct, non-overlapping patterns of regulation between segments. **D)** Schematic of GRN graph organization. **(E)** Cell type-specific GRNs define discrete regulation of T4 and L4 gene targets between glial populations.

We asked if transcription factors have inherently biased activity in glial cells, which depend on rostrocaudal positioning. To do so, we employed the Inferelator^44^ in multitask mode to construct gene regulatory networks (GRNs), creating a prior based upon transcriptionally active chromatin. Based on this analysis, we identified changes in TFA with cell type specificity across the thoracic and lumbar spinal segments. We assigned TFs based on their cognate recognition sequences, as provided by the JASPAR TF database^45^. We found that a subset of TFs are globally activated in a segment-dependent manner across multiple cell types, while others show strong cell type dependency (Figure 3B). None of the TFs we examined were inversely enriched between segments in different cell types. Of the globally enriched TFs, many have a role in modulating inflammatory responses in the CNS. In the lumbar region, astrocytes and OPCs have elevated activity of DDIT3, a key regulator of the unfolded protein response, which could position these cell types to transition more towards reactivity than their thoracic counterparts^46-48^. Oligodendrocytes and microglia show elevated NFKB1, which constitutes opposing effects in these cell types: protective against inflammation in oligodendrocytes^49^, and pro-inflammatory in microglia^50^. In the thoracic region, NFIA activity is enriched in astrocytes, OPCs, and oligodendrocytes, creating a differential potential to responses to injury or insult^51,52^. This analysis revealed that glial transcriptional programs are poised to differentially react depending on which segment of the spinal cord they reside, and could therefore contribute to the differential responses seen across segments under diseased conditions.

We then identified unique regulatory programs that define cell type-specific rostrocaudal differences in the spinal cord (Figure 3C, Supplementary Figure 3B, Supplementary Table 2). We found that lumbar and thoracic astrocytes have inverse activity of CPEB1 and EGR1, respectively, corresponding to phenotypically distinct populations: migratory^53^ vs neurotrophic^54^. OPCs show differential activity of key oncogenic transcription factors, with pro-tumor MYBL1^55^ and MYBL2^56^ overrepresented in the thoracic region and tumor-suppressing YY1^57^ and ZIC1^58^ in the lumbar region. This observation is consistent with the clinical consensus that a predominance of spinal cord tumors is located in the thoracic region of the spinal cord^2^. Relative to their thoracic counterparts, oligodendrocytes in the lumbar segment have increased baseline activity of TFs associated with stress responses traditionally seen in response to inflammation and remyelination, including ATF6A^59^, STAT3^60^, and IRF7^61^. Microglia in the thoracic segment enact homeostatic maintenance programs through the activity of TFs, including IKZF1^62^, ZNF768^63^, and ZNF306^64^, while lumbar microglia are more reliant on the activity E2F1^65^, VDR^66^, and SIX2^67^, suggesting a mechanism for how an imbalance in a seemingly global signaling program can trigger a region-specific response in microglia that then spreads to other regions. We built cell type-specific GRNs to understand how gene targets are affected by anatomically skewed TFAs (Figure 3D, E). While TFA enrichment can be shared by multiple cell types, we found that their gene targets are largely cell type-specific and non-overlapping between TFs (Supplementary Table 3). This observation suggests a finely tuned regulatory program in which a disturbance at a single node is propagated internally within a cell population before spreading to other cell types through a secondary mechanism.

### Dynamic Enhancer Activation Can Proceed in the Absence of Changes in Chromatin Accessibility

Having established that integrating chromatin accessibility and histone valence reveals novel regulatory strategies in a stable population of cells, we asked how dynamic remodeling of chromatin drives cell state transitions. We leveraged the continuous adult differentiation of OPCs to oligodendrocytes as a model for an active developmental trajectory. Oligodendrocyte populations are replenished at 0.3% per year in the healthy adult human CNS, a rate much lower than the mouse ^68,69^. Despite the importance of these adult-born oligodendrocytes in disease-associated remyelination^70^, the regulatory logic associated with adult gliogenic commitment is poorly understood. Furthermore, the molecular basis for the development of two oligodendrocyte subtypes has not been resolved.

We built a pseudotemporal trajectory to determine the time-ordered sequence of events that control the differentiation of OPCs into one of the two mature oligodendrocyte lineages. A bottleneck to studying branched differentiation trajectories is the difficulty in computationally modeling bifurcations in pseudo temporally ordered cells. Traditional approaches, such as RNA velocity^71^, are insufficient to model the transitions that OPCs undergo during differentiation (Figure 4A). To overcome this limitation, we used single cell Topological Data Analysis (scTDA), an algorithm that retains the shape of data in high dimensional space^72^. scTDA provides a continuous developmental trajectory, thus pseudotemporally ordering the cells with efficacy (Figure 4B). In contrast to RNA velocity, scTDA revealed three branches in the OPC to oligodendrocyte transition centered around an expectedly sparse population of actively differentiating precursors (Figure 4C, Supplementary Figure 4A). We assigned a starting pseudotime of 0 to cells in the root node, a position in the graph that maximizes both transcriptional entropy and, subsequently, the distance to terminal nodes in the graph. The root node resides within the actively differentiating OPC population. From this node, cells traverse one of 3 trajectories: they can transit from active to quiescent OPCs (Branch 0), commit to an Oligo1 cellular fate (Branch 1), or commit to an Oligo2 cellular fate (Branch 2). Cells in nodes proximal to the rooted node, as calculated by the minimal number of edges between them, are assigned an early pseudotime, while cells distal to it are assigned a late pseudotime. scTDA reveals that the discrete OPC clusters seen by UMAP using traditional dimensional reduction are smoothly distributed along a continuum of cell states, suggesting that the physiological functions of OPC subtypes are directly linked to their distance from terminal differentiation. Conversely, the less defined UMAP division between two oligodendrocyte subtypes resolved into a developmentally demarcated split between two distinct mature populations. Driving this branched fate decision is a change in the transcriptional profiles of cells in pseudotime (Figure 1D, Supplementary Figure 4B). We identified the regulatory program responsible for this divergent fate commitment by analyzing motif activity across pseudotime in each of the branches (Figure 4E). Like neonatal OPCs, adult OPCs are maintained in a quiescent state by the transcription factor SOX5^73^. These progenitors undergo a cascade of regulatory changes as they transition to active OPCs, driven by the activity of ASCL1, a transcription factor associated with remyelination programs^74^. FOXO1 is a regulator of oligodendrocyte differentiation in mice^75^, and is part of the regulatory progression that instigates Branch1 specification. NKX6-2, a transcription factor implicated in mouse oligodendrocyte differentiation^76,77^, has elevated activity selectively during Branch2 oligodendrocyte specification. These changes in motif activities across branches suggest a dynamic remodeling of chromatin and altered enhancer states that promote OPC to oligodendrocyte differentiation.

**Fig. 4.**
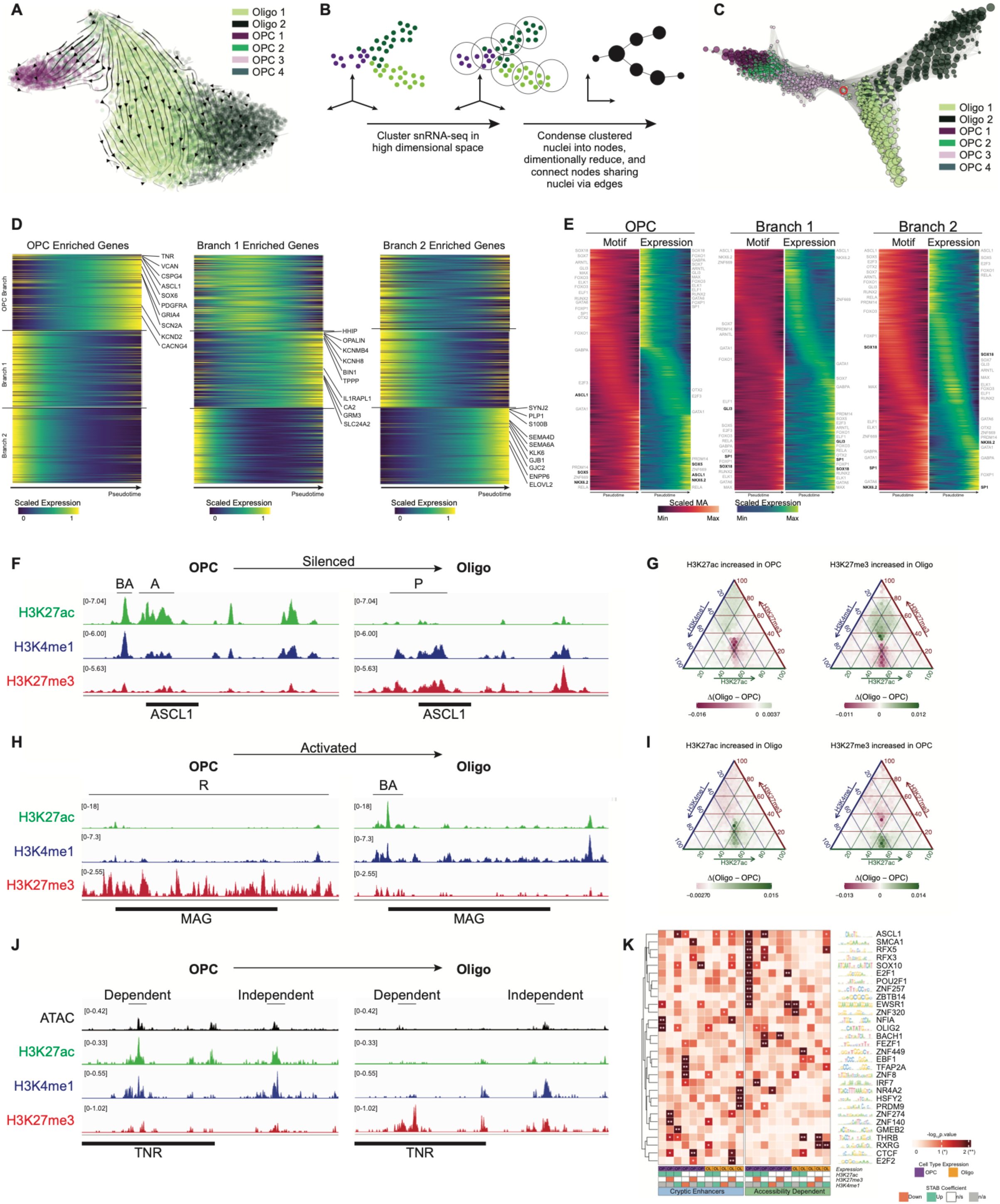
Gene Regulation of Adult OPC to Oligodendrocyte Differentiation. **(A)** Pseudotemporal ordering of OPCs and oligodendrocytes using RNA velocity does not recapitulate the trajectory of OPC to Oligodendrocyte differentiation. **(B)** Schematic of scTDA, an algorithm for pseudotemporal ordering of branched processes and graph-based analysis of differential gene expression and chromatin accessibility. **(C)** scTDA applied to the adult OPC to oligodendrocyte differentiation process reveals 3 branches in oligodendrocyte cell fate specification: OPC (Branch 0), Oligo1 (Branch 1), and Oligo2 (Branch 2). The rooted node (starting point pseudotime 0) is centered in the graph and outlined in red and falls within the actively differentiating OPC population. **(D)** Scaled expression of genes enriched in a single branch, plotted as a function of pseudotime across the three branches, shows minimal overlap in temporal expression patterns with other branches. **(E)** Independently pseudotemporally ordered motif activity (red) and transcription factor expression (green) for each branch shows a wave of regulatory logic associated with cell fate determination. **(F)** Silencing of ASCL1 gene expression in mature oligodendrocytes corresponds to a regulatory shift at enhancer loci from bivalent active (BA) and active (A) to poised (P) states. **(G)** Triangle plot demonstrating consistent inverse relationships between H3K27ac and H3K27me3 signal at enhancer peaks that are active in OPCs (left) and silenced in oligodendrocytes (right). **(H)** The MAG locus, a myelinating gene expressed upon oligodendrocyte differentiation, is fully repressed in OPCs (R), and is derepressed and gains a bivalent active (BA) enhancer peak upon differentiation. **(I)** Triangle plot demonstrating consistent inverse relationships between H3K27ac and H3K27me3 signal at enhancer peaks that are active in oligodendrocytes (left) and silenced in OPCs (right). **(J)** The TNR locus shows a complex regulatory pattern in which two peaks, both bivalent active in OPCs and poised in oligodendrocytes, show differential dependence on chromatin accessibility changes. The intronic regulatory peak is silenced in concordance with the loss of chromatin accessibility, while the upstream peak changes chromatin valence independently of chromatin accessibility change. **(K)** Matrix plot showing the significance of TFA scores between OPCs and oligodendrocytes organized by TF preference for cryptic vs. remodeling-dependent enhancers between cell types. Peak-associated gene expression was calculated to be enriched in OPCs (OP) or oligodendrocytes (OL). The STAB coefficient, defined as the level of modification enrichment at TF-associated peaks between the two cell types, was calculated as enriched (up), depleted (down), not significant (n/s) or not considered (n/a). TFs demonstrate largely mutually exclusive preferences for valence at enhancer regions that either correlate with changes in chromatin accessibility (accessibility dependent) or are independent of chromatin accessibility changes (cryptic enhancers).

We then examined changes in histone modifications at regulatory sites during oligodendrocyte differentiation. ASCL1, a transcription factor that is active specifically in OPCs ^74^, shows a combination of two enhancer states in progenitor cells (active and bivalent active). Upon repression in mature oligodendrocytes, both sites transition to a poised, but not fully silenced, state (Figure 4F). This is a reproducible shift across enhancers for genes that are repressed upon cell fate commitment (Figure 4G). Conversely, the genomic region encompassing the proximal regulatory and coding sequences for the mature oligodendrocyte myelination gene MAG is fully repressed in OPCs. Upon differentiation, the enhancer for MAG becomes bivalent active while the TSS region is fully active, supporting a de-repression model for genes activated in mature oligodendrocytes (Figure 4H). These dynamics are reproducible across distal enhancers for genes that are induced upon oligodendrocyte differentiation (Figure 4I).

Finally, we asked whether these enhancer dynamics occur dependently or independently of chromatin accessibility changes. Two enhancers of TNR, a gene expressed in OPCs, illustrate the complexity of this regulatory mechanism. The upstream enhancer switches from bivalent active to poised independent of chromatin accessibility changes, while the intronic enhancer is silenced and concordantly decreases in accessibility (Figure 4J). A comprehensive analysis of enhancer peaks involved in the OPC to oligodendrocyte differentiation reveals almost mutually exclusive regulatory dynamics of TFs that rely on chromatin accessibility changes and those that do not (Figure 4K). OLIG2, the critical developmental bHLH transcription factor, serves as a master regulator of oligodendrocyte differentiation and a crucial activator of myelination genes and has an integral impact on glioblastoma and responses to injury and disease^78-80^. Despite OLIG2 motif activity being a major determinant of OPC identity when considering glial cell heterogeneity in ATAC measurements, a striking property of the OLIG2 consensus sequence is its localization in cryptic enhancers. The activity of OLIG2 is enriched in OPC-specific genes at constitutively accessible enhancers, where it is preferentially regulated by histone acetylation. Conversely, SOX10, the principal regulator driving oligodendrocyte differentiation, exhibits motif activity that acts in concert with changes in chromatin accessibility. Consistent with the de-repression model for mature oligodendrocyte gene activation (such as MAG), OLIG2 target expression in oligodendrocytes is inversely related to histone methylation. These results point to a complex regulatory program driving the formation and maintenance of adult oligodendrocytes in the spinal cord that depends in equal parts on chromatin restructuring and remodeling of histone valence at constitutively accessible sites.

### Spatially Organized Cellular Networks in the Human Spinal Cord

Neuronal cytoarchitecture is highly stereotyped in the spinal cord, with motor columns and Rexed laminae defining neuraxial positioning. The corresponding glial organization is not well defined. Given the importance of proximal paracrine signaling in mediating intercellular communication, we asked how astrocytes, microglia, OPCs, and oligodendrocytes pattern across the cross-section of the spinal cord to facilitate homeostatic function. Specifically, we asked if glial cells form local cellular *networks* that are distinct from traditional cytoarchitectural constraints and have, therefore, been overlooked. We examined the spatial organization of cells in the L4 lumbar segment of a deceased transplant organ donor, developing an approach to identify and quantify patterns of stereotypical cellular networks along the dorsoventral and mediolateral axes (Figure 5A). We used STARmap^81^ to spatially profile 146 genes with single cell and single molecule resolution, inclusive of glial subtype-specific markers (Supplementary Table 4). We clustered the *in situ* profiled cells and observed a reproducible transcriptional signature of glial subtypes consistent with the snRNA-seq dataset (Figure 5B, Supplementary Figure 5A) and generated a cartograph to spatially identify cell types in the tissue (Figure 5C). For each cell, we calculated, within a radius of 60µm, the composition of the surrounding cell types. These proximal cells formed the basis for calculating a cellular network (CN). We defined the network profile of each cell as the cumulative count of cell types within the given radius. We aggregated the neighborhood profiles for each cell type by performing k-Nearest Neighbor (k-NN) clustering^82^ and determined cluster stability by bootstrapping. Community detection was performed on the resulting k-NN graphs to identify repeated cellular networks tiling across the lumbar segment (Figure 5D, Supplementary Figure 5B-F).

**Fig. 5.**
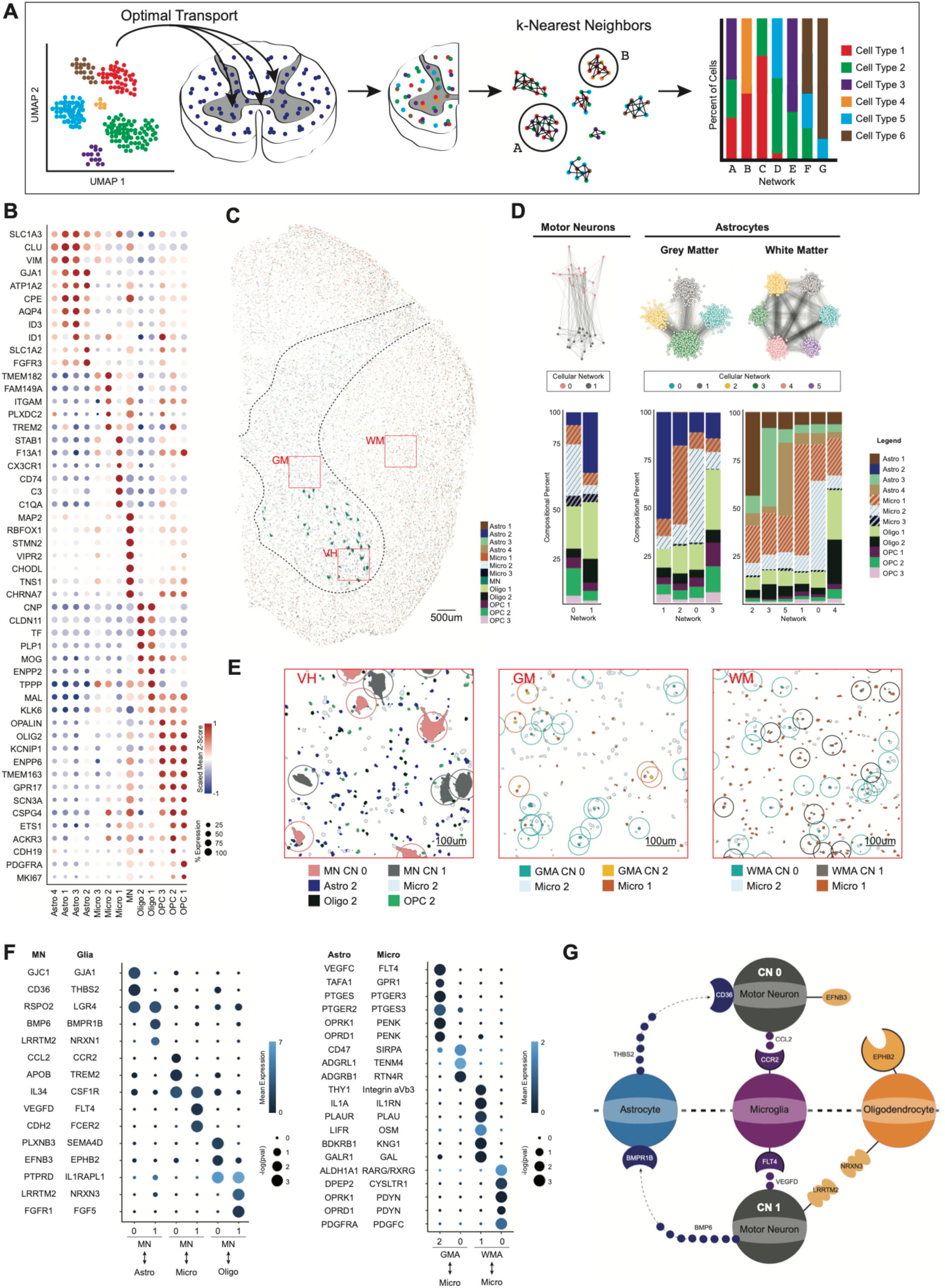
Spatial Transcriptomics and Community Detection Identify Stereotyped Neighborhoods of Cellular Composition. **(A)** Schematic overview of spatial data analysis and community detection. Optimal transport is used to accurately project high-depth snRNA-seq measurements onto *in situ* profiled cells. For each cell type, the spatial relationships between cells are calculated by kNN clustering, and cellular networks are defined by reproducible and stable communities of proximal cells. **(B)** Dot plot of *in situ* transcriptomic data identifies glial clusters consistent with snRNA-seq based on scaled expression of marker genes. **(C)** Cartograph showing glial subtypes and MNs in the *in situ* sequenced L4 spinal cord cross-section. The dotted black line designates the grey matter/white matter boundary. The red boxes correspond to representative regions of white matter (WM), grey matter (GM), and ventral horn (VH). **(D)** Top: kNN graph and community detection identify cellular networks (CNs) for MNs (left), grey matter astrocytes (center), and white matter astrocytes (right). Bottom: stacked bar graphs show the cumulative percent contribution of proximal cell types for each CN. **(E)** Zoomed in cartographs of boxed regions in C visualizing enriched cell types in two CNs for MNs (left), GMA (middle), and WMA (right). For clarily, only the cell types that are differentially enriched between CNs are shown. For each plot, the analyzed cell type is outlined by a circle (MN radius 80µm, GMA and WMA radius 60µm), within which cells are considered proximal. Cell type stoichiometry within the radius for each community is reproducible, tiles across the tissue, and is consistent with the stacked bar graph in D. **(F)** CellphoneDB analysis highlighting ligand-receptor pairs that are unique to cell type interactions within a CN. Left: MNs in CN 0 and 1 show distinct interactions with astrocytes, microglia, and oligodendrocytes. Right: GMAs in CN 0 and 2 and WMAs in CN 0 and 1 show distinct interactions with microglia. **(G)** Schematic of different signaling pathways that MNs in CN 0 and CN 1 participate in.

CNs define a reproducible stoichiometry of cells proximal to the cell type being analyzed. When considering motor neurons, we discovered that they segregate into two networks: Network 0, in which motor neurons are surrounded predominantly by astrocytes (GJB6^+^), and Network 1, in which motor neurons are surrounded predominantly by Micro2, a microglial population enriched for P2RY12^+^, a purinergic receptor that characterizes motile microglia. These findings strongly point to preset vulnerability and resistance in motor neurons that may be linked to pre-existing intercellular cues. We also observed that Network 0 motor neurons have significantly more proximal Oligo2 (KLK6^+^) neighbors, while Network 1 has more OPC2 (MET^+^) neighbors. These observations provide a rationale for differential paracrine signaling that may confer resistance or vulnerability to motor neuron stress. In contrast to motor neurons, more recent studies indicate that key differences in white matter and grey matter astrocytic reactivity occur during both aging and neurodegenerative disease. As a reflection of their diverse functional impact on neuronal homeostasis, astrocytes display regional and key molecular differences. We, therefore, sought to extend our analyses to capture the impact of cellular networks on the astrocytic state. Grey matter astrocytes (Astro2, GJB6^+^) are found in 4 CNs: Network 0 astrocytes are commonly found near Micro2 (P2RY12^+^) microglia, and Network 2 astrocytes preferentially associate next to Micro1 (SPP1^+^) microglia. Network 1 astrocytes localize adjacent to other grey matter astrocytes, and Network 3 astrocytes reside proximal to Oligo1 (OPALIN^+^). In the white matter, Astro1 (AQP4^+^) reside in 6 CNs, preferentially neighbored by Micro2, Micro1, Astro1, Astro3 (CNTNAP1^+^), oligodendrocytes, or Astro4 (RFX4^+^). These CNs are tiled across the spinal cord and are spatially intermingled (Figure 5E).

To understand the functional significance of the organization of these CNs, we characterized the intercellular signaling pathways within individual neighborhoods of cells. We used optimal transport to project high-depth gene expression measurements from snRNA-sequencing onto our spatially profiled data. This provided a spatially resolved dataset with transcriptional depth. We then used CellphoneDB^83^ to identify receptor-ligand interactions between distinct cell types within each CN. We identified a panel of unique receptor-ligand pairs that are dedicated to cells in different spatial communities (Figure 5F, Supplementary Table 5). The information exchanged between grey matter astrocytes and microglia is different depending on which community they are a part of and distinct from the information exchanged between white matter astrocytes and microglia. These spatially restricted patterns of signaling reveal a previously unrecognized level of cellular heterogeneity and provide additional insights into the selective vulnerability of cells to stress, inflammation, and neurodegeneration (Figure 5G).

## DISCUSSION

In this study, we accomplished four principal objectives. First, we established a multiomic cell atlas of the thoracic and lumbar segments of the healthy adult human spinal cord, defined the corresponding cis-regulatory elements driving their specification, and established an optimal transport approach to integrating these data. Second, we developed and applied STAB-seq to track enhancer states and chromatin remodeling in individual spinal cord nuclei. We uncovered previously unidentified cryptic enhancer classes, defined their dynamics in both stable cellular populations and in actively differentiating cells, and proposed a potential role for enhancer chromatin valence in disease processes. Cryptic enhancer transitions challenge multi-omic studies predicated upon chromatin potential as the arbiter of gene activation and bridge the dissonance commonly observed between mRNA expression and ATAC-inferred gene activity. Third, we defined anatomically localized transcription factor activities and concordant reorganization of cell type-specific gene regulatory networks across the rostrocaudal axis of the spinal cord. We extended this approach to identifying the regulatory dynamics of oligodendrocyte subtype specification and transition to quiescence in OPCs. Finally, we demonstrated that cellular identity can be recast in the context of cellular networks. We also described network-specific paracrine signaling pathways based on expressed receptor-ligand pairs, which support homeostasis. Critical in these findings is the redefinition of alpha motor neurons by the cellular neighborhoods in which they reside, characterized by distinct distributions of proximal astrocytes, microglia, and oligodendrocyte subtypes. In the white matter of the spinal cord, two populations of seemingly identical astrocytes are selectively engaged by phagocytotic or scavenging microglial populations. Such repeatable and stereotyped cellular neighborhoods provide evidence of a previously unappreciated cytoarchitectural axis in the spinal cord.

### Nonequivalent Regulatory Potential Across Spinal Segments

Previous studies of the mammalian spinal cord focused on transcription as an arbiter of cellular diversity and function^8^. We extend those results, profiling histone modifications and transcriptional state in spinal nuclei isolated from the thoracic and lumbar regions of the spinal cord. The data revealed distinct differences in transcription factor activities between glial subpopulations along these anatomic axes. More broadly, our studies identified substantial rewiring of glial gene regulatory networks along the rostrocaudal axis. Defined as the influence of transcription factors (TFAs) on gene activation, differences in TFAs within glial subpopulations suggest critical differences in the potential responsiveness of these cell types to stress. Disease Associated Glia (DAGs) in the spinal cord have been well-documented in the context of transcriptional readouts, with perplexing focal pathological effects^84^. Our work suggests that DAGs may result from seemingly identical glial cells, from a transcriptional vantage point, with differing underlying transcription factor activity. During neurodegenerative disease conditions, such as amyotrophic lateral sclerosis, disease initiation is asymmetric. By way of example, thoracic onset ALS is exceedingly rare, impacting 3% of cases^85^. Our work points towards the need for further study of glial contributions to selective motor neuron degeneration in context of the anatomic influence transcription factor families have on transcription.

### Cryptic Human Enhancers Impact Regulatory Dynamics

Chromatin potential, defined as the reorganization of chromatin towards accessibility to transcription, has been described as a predictor of transcriptional activation. Traditional single nuclei multiomic assays rely on correlations between changing chromatin accessibility and RNA abundance, and therefore depend upon chromatin potential for regulatory site determination. We hypothesized that cryptic enhancers exist that activate transcription, absent canonical chromatin potential. We, therefore, developed and applied STAB-seq with a specific interest in detecting enhancers within the thoracic and lumbar segments of the spinal cord that could refine our understanding of glial gene activation. We identified cryptic enhancers, controlling transcriptional activation absent discernible reorganization of chromatin accessibility in both static glia and differentiating oligodendrocyte progenitor cells. These enhancers are best defined as upstream regulatory regions that are constitutively accessible, and are identified through transitions towards H3K27Ac modification and subsequent gene activation. We identified these regulatory regions across all major glial subpopulations, demonstrating their ability to define glial subtypes, their potential contributions towards cellular reactivity, and their involvement in oligodendrocyte subtype specification. The constitutive accessibility of these enhancers may arise from several biological processes. On one hand, these enhancers may arise from developmental considerations. For example, progenitors such as pMNs, that give rise to both OPCs and motor neurons^86^, may yield mutually exclusive cryptic enhancer states in each cell type. A developmental dead end in an OPC may result in a constitutively accessible state that is activated in motor neurons. Alternatively, the accessibility of these enhancers may result from mitotic bookmarking during development, whereby transcription factor binding during mitosis mitigates chromatin closure and enhances TF binding to its cognate binding site in daughter cells^87^. Interestingly, the presence of these enhancers may increase in the brain and spinal cord as a function of age. Studies of epithelial stem cells have demonstrated that inflammation in these cells renders their daughter cells poised to activate immune genes through maintenance of chromatin accessibility^88^. We reason that stressors occurring during aging may increase the abundance of these cryptic enhancers, rendering glia and neurons poised to activate responsive transcriptional programs. Ultimately, our studies point towards the importance of transcription factor binding as a more faithful indication of transcriptional activity rather than chromatin accessibility, which has been argued elsewhere^89^. Both the impact of these cell type specific cryptic enhancers on the kinetics of gene activation and their intersection with noncoding genetic variation in neurodegeneration requires further investigation.

### Cytoarchitectural Organization as a Framework for Cellular Identity

We hypothesized that, rather than being randomly distributed throughout the spinal cord, each glial cell resides within one of several stereotyped intercellular networks. If true, cellular identity could be recast in the context of surrounding cells. We applied the Starmap *in situ* sequencing approach for clarification, and developed a computational framework using community detection to test our hypothesis. Previous studies in the mouse cortex showed that non-neuronal cells become spatially proximal during aging, preferentially colocalizing, pairwise, with cell type specificity^90^. Our studies demonstrate that glia not only have a pairwise preference for proximal cells, but they form repeated cellular networks that organize across the tissue. Importantly, our analysis demonstrated that alpha motor neurons appear to participate in one of two networks, preferentialy surrounded by different distributions of proximal astrocytes, microglia, and oligodendrocyte subtypes. Historically, differences between motor neuron subclasses have relied upon intrinsic transcriptional definitions that struggle to explain selective resistance or vulnerability to degeneration. One possibility is that selective motor neuron vulnerability in ALS is a consequence of vulnerable motor neuron populations within in the observed networks enriched for astrocytes, aggravating local signalling that has been shown to impact neural viability^91^. A bulk spatial study of the human spinal cord in ALS has shown that disease severity correlates with proximity to the initial site of symptom onset, however this study did not have the spatial resolution necessary to define cell type-specific contributions to disease^92^. White matter and grey matter astrocytes, known to occupy non-overlapping spatial territories that have traditionally been considered self-contained^93^, also reside within discrete cellular networks with stereotyped neighboring cells. We postulate that these networks facilitate dedicated paracrine signaling, and identify the expression of unique receptor-ligand combinations between astrocyte, microglia, and oligodendrocyte subtypes that reside in different networks. Our work reveals an enrichment of phagocytic microglia within a white matter astrocyte network, suggesting that cytokines produced by these microglia have the potential to drive reactive gliosis and elicit a focus for neurodegeneration^94^.

Taken together, the approaches developed in this study reveal multiple layers of spatial and epigenetic regulation of cell states in the healthy human spinal cord. Our findings invite a deeper exploration of the impact of local intercellular communication and chromatin remodeling on the diverse cellular transitions observed during neurodegenerative disease. Although an abundance of resources exists to define heterogeneity within the central nervous system, our work challenges the notion that the transcriptome is sufficient to capture cellular identity. Rather, poised enhancer states and local cellular networks providing inductive paracrine signals provide a deeper insight into cellular populations predisposed to resilience or degeneration during injury or insult in the human spinal cord. This approach, applied to parallel studies of diseased states is likely to provide novel insights into neurodegenerative disease mechanisms.

#### Limitations of the study

This study offers multiomic and single cell resolved spatial transcriptomic data generated from tissue isolated from deceased transplant organ donor non-neurological control cases. There does not exist a published dataset for comparison and therefore our study is statistically underpowered with respect to small or rare cellular subpopulations. We therefore have not included these cellular groups in our analyses. Given that single cell spatial transcriptomic approaches, such as Starmap, require probe panel design and therefore there may be additional transcriptional events that characterize intercellular interactions.

#### Lead contact

Further information and requests for resources and reagents should be directed to and will be fulfilled by the lead contact, Abbas Rizvi.

## ACKNOWLEDGMENTS

We thank Neil Shneider for contributing post-mortem tissue. All authors thank the donors and their families for granting access to post-mortem and deceased transplant organ donor tissue. This study was supported by the Chan Zuckerberg Initiative (2017-174051, 2018-190766 & RG98793) and the Department of Defense (W81XWH-22-1-0114). The work of R.R., A.W., and J.F. was supported by NIH (R35CA253126) and NSF (#1912194).

## AUTHOR CONTRIBUTIONS

Experimental Design, E.K.K, A.H.R., T.M.; STAB-seq Method Development, E.K.K., A.H.R; Optimal Transport, M.C., J.L., A.W., E.K.K. A.H.R.; Bioinformatics, A.W.; STAB-seq Analysis, W.L., A.W, E.K.K, A.H.R; Starmap Experimentation and Image Processing, L.P., E.K.K, A.H.R.; Community Detection, A.W., M.C., E.K.K., A.H.R.; Gene Regulatory Network Construction, A.T., W.L., Y.X., scTDA, J.F., Manuscript Preparation, E.K.K. A.H.R., T.M.; Supplementary Methods and Manuscript Review, All Authors; Funding Acquisition, A.H.R. T.M. and R.R.; Reagent Preparation, W.P.; Deceased Transplant Organ Donor Tissue Acquisition K.T.M. and K.S.P.

## SUPPLEMENTAL FIGURES

**Supplementary Figure 1:**
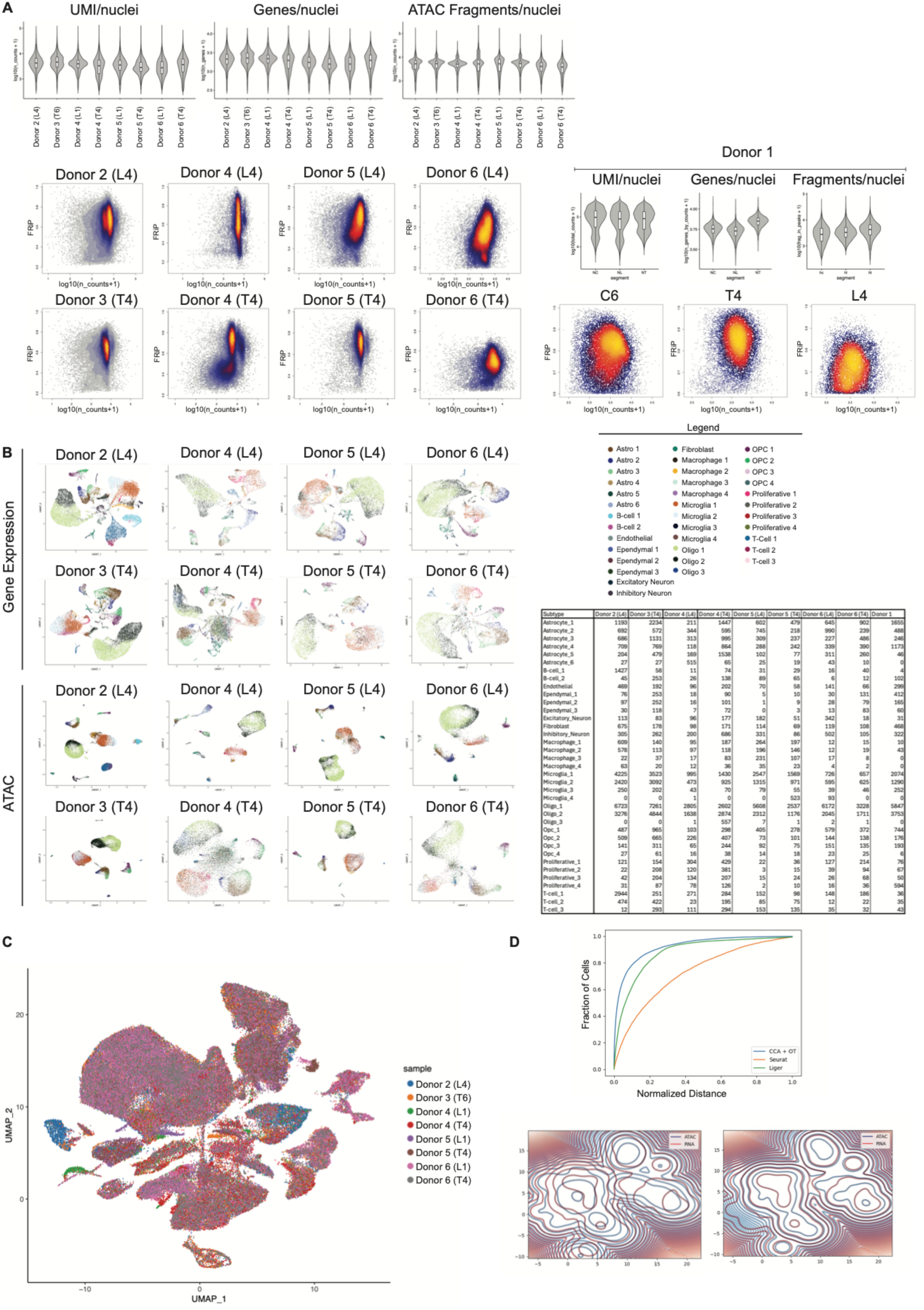
Libraries across donors show consistent quality control metrics. A) Unique molecular identifiers (UMIs), genes detected, ATAC fragments, and fraction of reads in peaks (FRiP) are consistent across all deceased transplant organ donor tissues and between segments for donor 1 (analyzed separately). B) RNA and ATAC UMAP representations for each sample show consistent cellular heterogeneity. C) Batch effects are not observed when merging libraries from deceased transplant organ donor tissues. D) OT outperforms other methods for co-embedding RNA and ATAC measurements into a shared feature space. Top: ROC for OT (blue), Liger (green), and Seurat (orange). Bottom: contour plots show enhanced coembedding of RNA and ATAC datasets using OT (right) compared with Seurat (left).

**Supplementary Figure 2:**
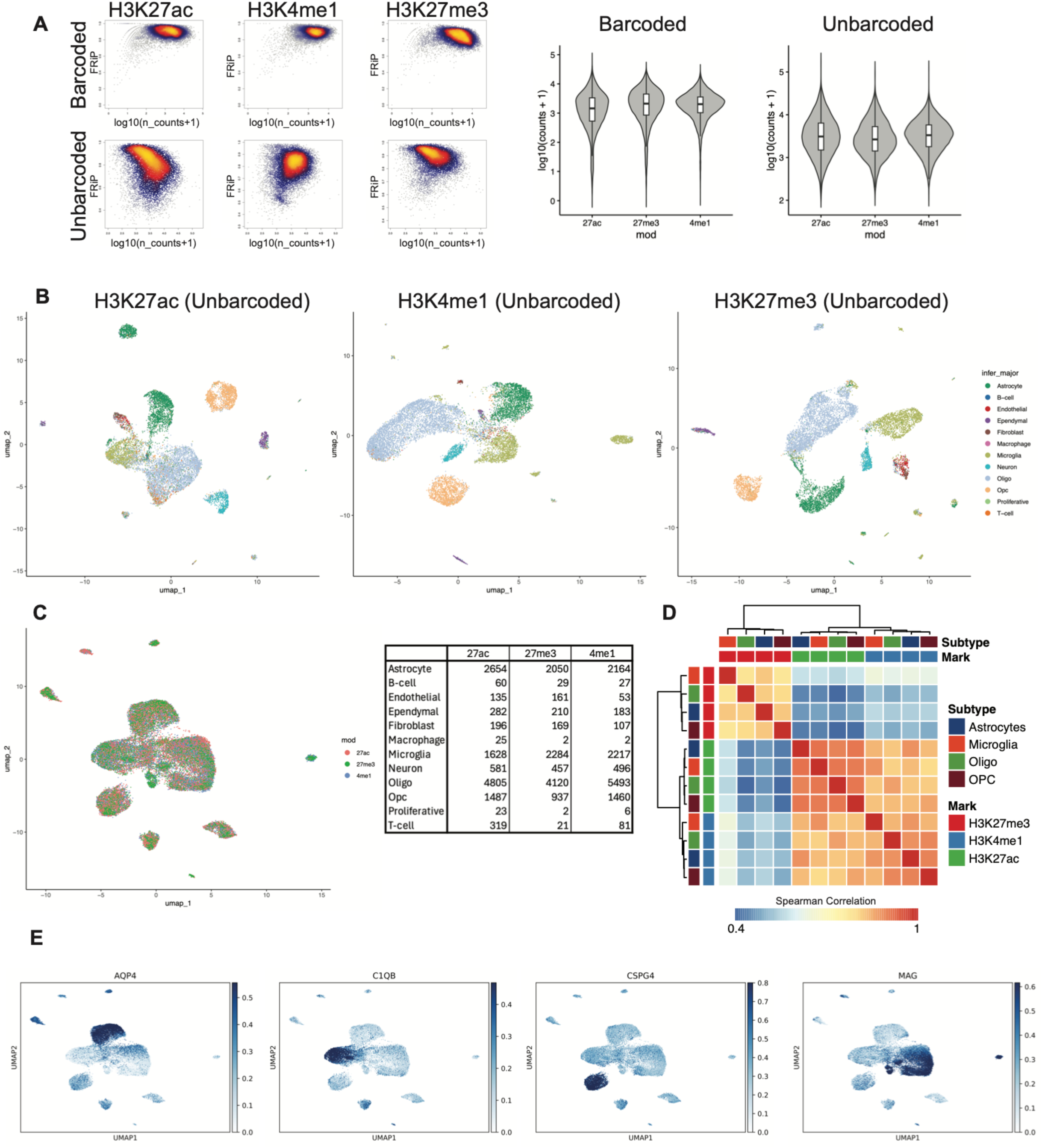
STAB-seq integration of gene expression with chromatin valence. A) Barcoded (dual calling card, modification specific) and unbarcoded (global Tn5 tagmentation product) FRiP metrics and fragment counts per nuclei are consistent between the three modifications. B) UMAP representations of STAB-seq libraries show consistent major cell type recovery between modifications profiled. C) Coembedding the unbarcoded reads from all STAB-seq samples show no batch effects in the UMAP representations and consistent cell type recovery compared with traditional ATAC profiling. D) Dual calling card containing fragments show expectedly low spearman correlation between silencing (H3K27me3) and activating (H3K27ac) histone marks between cell types. E) OT pairing of multiome sequencing with STAB-seq enables accurate integration of gene expression with STAB-seq histone modification in single cells, as shown by STAB-seq UMAP representations colored by inferred RNA expression.

**Supplementary Figure 3:**
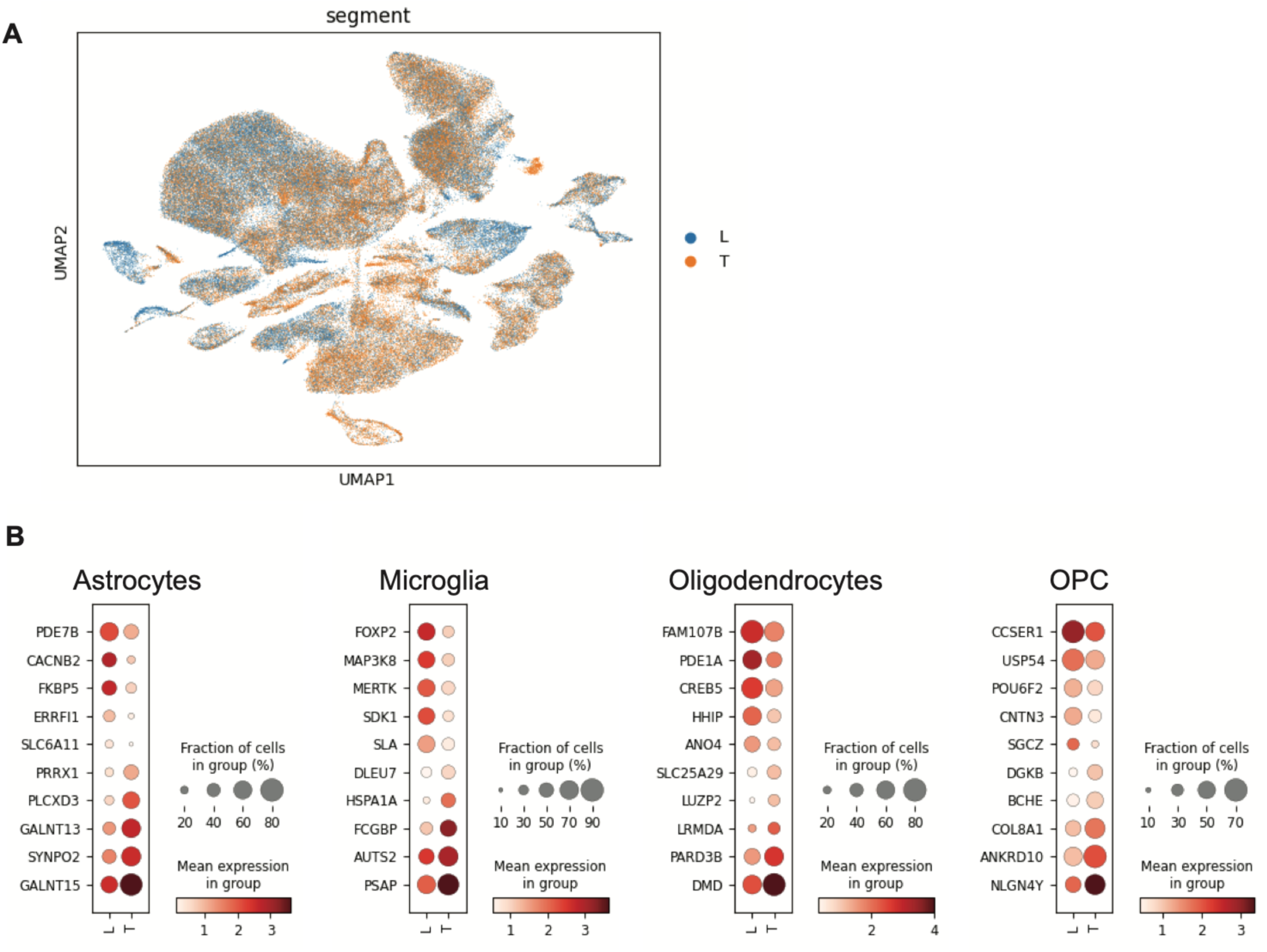
Segment dependent gene expression differences between the thoracic and lumbar spinal cord. A) UMAP representation of thoracic and lumbar nuclei demonstrates that segment level differences cannot be resolved through traditional clustering methods based on gene expression. B) Dotplots demonstrating cell type specific gene expression differences between two spinal cord segments.

**Supplementary Figure 4:**
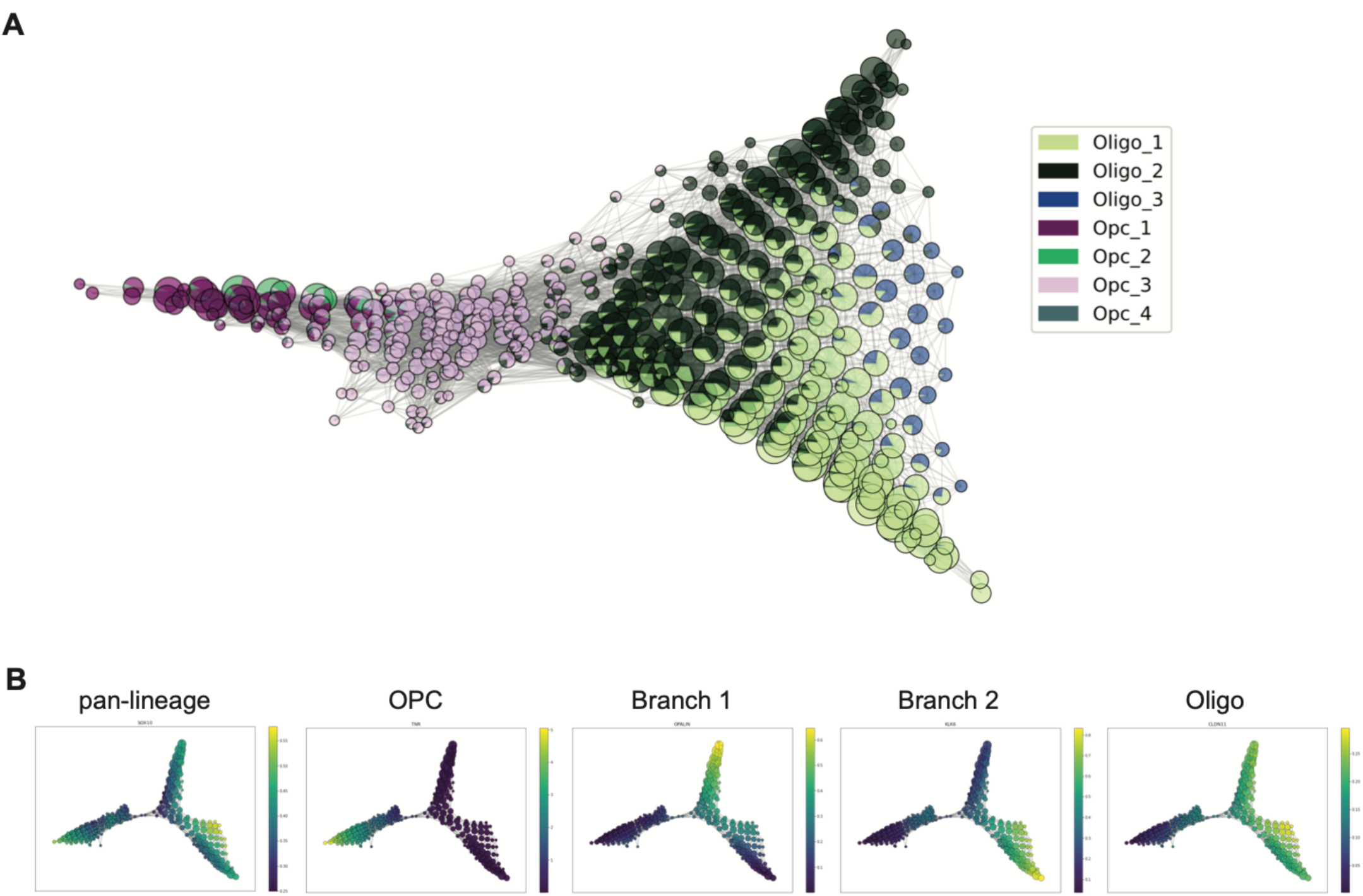
scTDA representations of the OPC to oligodendrocyte differentiation trajectory. A) The scTDA representation is consistent between Donors 2-6 (shown) and Donor 1. B) Scaled gene expression of subtype-specific genes along the scTDA trajectory shows enrichment between branches and throughout differentiation.

**Supplementary Figure 5:**
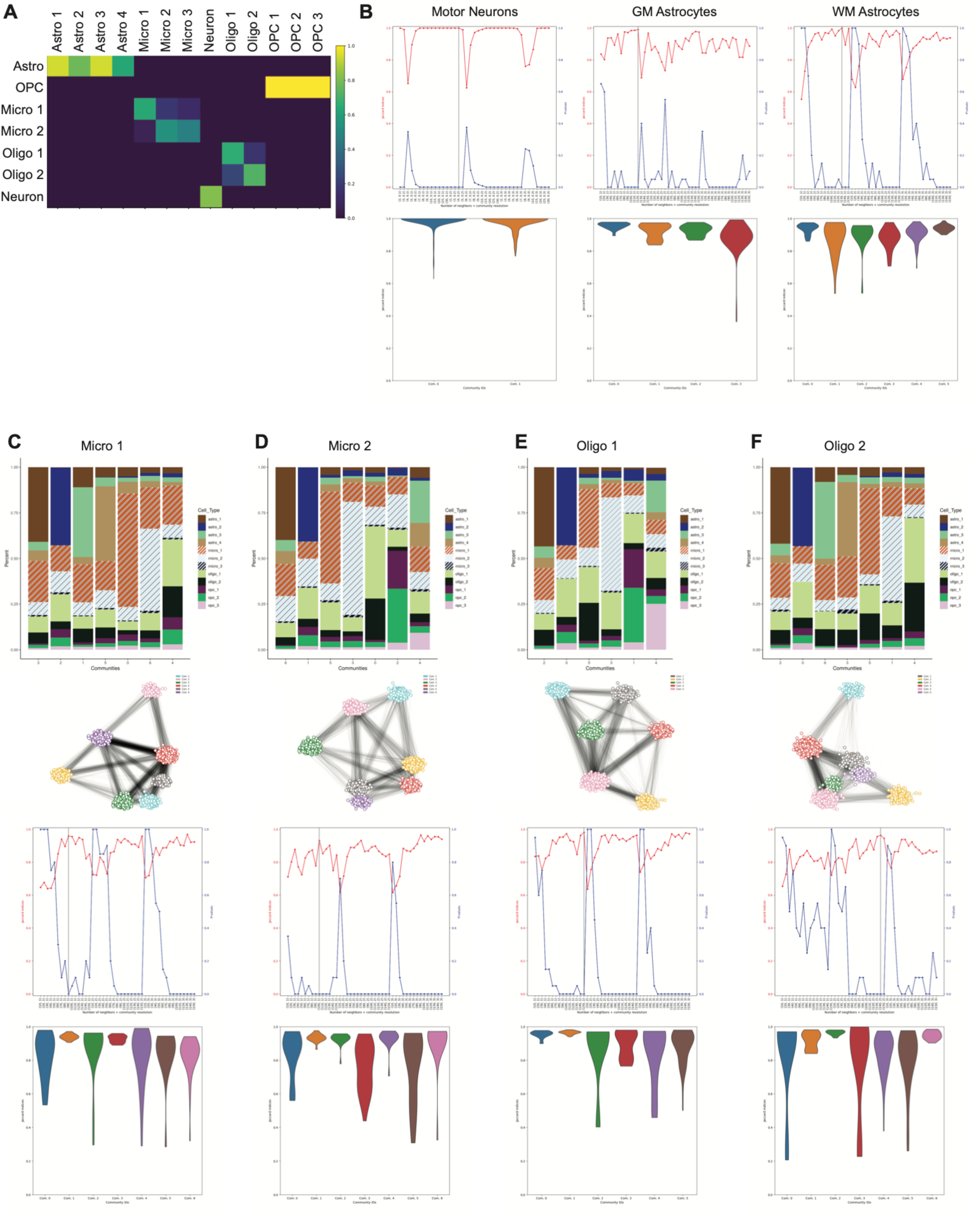
Stereotyped Cellular Networks between glial subtypes in the spinal cord. A) OT calculated confusion matrix between nuclei in clusters profiled via 10x and the nuclei in clusters defined through STARmap show strong consistency in cell type identification (scaled by number of nuclei in each cluster). B) Stability of cellular networks for MNs, GMA, and WMA. Top: the resolution parameters (grey vertical line) for each cell type were chosen as the intersection of the highest jaccard index (red) and lowest p-value (blue). Bottom: violin plots showing stability of cellular networks through bootstrapping. C-F) Cellular networks for Micro1, Micro 2, Oligo1, Oligo2 in STARmap profiled spinal cord cross-sections. Top: Aggregated cell type contributions for CNs. Middle top: knn graphs for CNs for each cell type. Middle bottom: resolution parameters chosen via jaccard stability and p-value. Bottom: violin plots showing stability of CNs through bootstrapping.

## METHODS

### Study Participant Details

For five adult male, non-neurological control deceased transplant organ donors (Donors 2-6), tissues were acquired by the Collaborative Biorepository for Translational Medicine, from deceased transplant organ donors under Research Ethics Committee approval (ref 15/EE/0152, East of England Cambridge South Research Ethics Committee) and informed consent from the donor families. The spinal cord from one adult male donor (Donor 1) was resected post-mortem at Columbia University Medical Center under Research Ethics Committee approval and with informed consent from the donor family.

### Tissue Resection from Deceased Transplant Organ Donors

Samples were collected from donors proceeding to organ donation shortly after cessation of circulation; the chest is opened, the aorta is cross-clamped and organs are perfused under pressure *in-situ* with cold organ preservation solution (Belzer UW^®^ Cold Storage Solution, Bridge to Life (Europe) Ltd.) and cooled with topical application of ice. After the organs for transplantation were removed, the spinal cord samples were collected by removing a wedge of vertebrae from the T4 and L4 regions, exposing the vetebral foramen and dissecting a full thickness length of the corresponding region of the spinal cord. Samples for this study were all obtained within 60 minutes of cessation of circulation, placed in cold preservation for transport on ice (at 4°C) to the laboratory. The tissue samples were immediately padded dry with sterile filter paper and snap frozen in liquid nitrogen vapour on a Parafilm M (PM 999, Bemis Co Inc, Neenah, WI) coated aluminium foil boat before being stored at -80°C and transported on dry ice.

### Nuclei Isolation

50-100mg of tissue was shaved from a cross-section of the spinal cord on dry ice, transferred to 4mL of ice-cold homogenization buffer (5mM CaCl2, 3mM Mg(CH3COO)2, 10mM Tris-HCl pH7.8, 1mM DTT, 320mM sucrose, 0.1mM EDTA, 0.1% NP-40, 0.1mM PMSF) in a dounce homogenizer, and incubated on ice for 2 minutes. The tissue was physically dissociated using sequential 10 strokes of the loose pestle followed by 10 strokes of the tight pestle. The nuclei suspension was filtered through a 40um mesh filter into a 15mL conical tube, and diluted with an equal volume of ice-cold 50% Optiprep salt solution (50% Optiprep (Sigma, D1556), 5mM CaCl2, 3mM Mg(CH3COO)2, 10mM Tris-HCl pH7.8, 1mM DTT). The tube was gently inverted to mix. The resulting 25% Optiprep/nuclei suspension was layered over an isosmotic 29% Optiprep solution (29% Optiprep, 5mM CaCl2, 3mM Mg(CH3COO)2, 10mM Tris-HCl pH7.8, 1mM DTT, 160mM sucrose), and spun in a swinging bucket centrifuge at 6,000g for 30 minutes at 4C. The supernatant was completely removed through slurping off from the top of the meniscus to prevent debris carryover, and nuclei were resuspended in the appropriate buffer for downstream processing.

### Library Generation

#### 10X Multiome GEX+ATAC

The Chromium Next GEM Single Cell Multiome ATAC + Gene Expression reagents (10x Genomics, 1000285) were used to generate simultaneous cDNA and ATAC libraries. Briefly, nuclei were resuspended in 1mL Lysis Buffer (10mM Tris-HCl pH7.4, 10mM NaCl, 3mM MgCl2, 1% BSA, 0.1% Tween-20, 0.1% NP-40, 0.01% Digitonin, 1mM DTT, 1U/uL Protector RNase Inhibitor [Sigma-Aldrich, 03335402001]) and incubated on ice for 2 minutes, diluted with 1mL Lysis Buffer without detergents, and pelleted at 500g for 5 minutes. Nuclei were resuspended in 1x Nuclei Dilution Buffer + 1mM DTT + 1u/uL Protector RNase Inhibitor at 3,230 nuclei/uL, and assessed for structural integrity and monodispursion. 16,100 nuclei were input into 10x Genomics tagmentation followed by Chromium (Next GEM Chip J) droplet encapsulation. 7 cycles of pre-amplification PCR were performed, followed by 14 PCR cycles for gene expression (GEX) libraries and 7 PCR cycles for ATAC libraries. Libraries were purified with double size selection (GEX: 0.6x and 0.8x; ATAC: 0.6x and 1.25x) using SPRIselect (Beckman Coulter, B23317), and run on a Bioanalyzer for quantification and structure assessment. GEX libraries were multiplexed for sequencing on a NextSeq 550 using High Output 150 reagents at 28x90x10x10 cycles. ATAC libraries were multiplexed for sequencing on a NextSeq 550 using High Output 150 reagents at 50x49x8x16 with a custom sequencing recipe of 8 dark cycles for Index 2.

#### snATAC-seq

The BioRad SureCell ATAC-seq Library Prep Kit (BioRad, 17004620) was used to generate snATAC-seq libraries following manufacturer’s recommendation. Briefly, nuclei were resuspended in PBS + 0.1% BSA + 0.01% Tween-20 + 1x Roche EDTA-free protease inhibitor. Nuclei were concentrated to 3,640 nuclei/uL in PBS + 0.1% BSA, and assessed for structural integrity and monodispursion. 60,000 nuclei were input into SureCell tagmentation followed by ddSeq droplet encapsulation. 8 cycles of in-droplet indexing PCR was performed, and 2.5uL of the amplified product was input into KAPA (Roche, KK2602) qPCR to determine second-round PCR cycle number. Libraries were amplified to log-linear phase (7-9 cycles) with the SureCell ATAC-seq Library Prep Kit reagents, purified with two rounds of 1x AMPure XP (Beckman Coulter, A63881), and run on a Bioanalyzer for quantification and structure assessment. Libraries were multiplexed for sequencing on a NextSeq 550 using High Output 150 reagents at 118x40x8 cycles, with the custom Read 1 sequencing primer (BioRad SureCell ATAC-seq Library Prep Kit).

#### High Depth snRNA-seq

Nuclei were resuspended in PBS + 0.2U/uL Superasin (Thermo Fisher, AM2694) + 1ug/mL DAPI. Single nuclei were sorted on a high speed MoFlo XDP FACS sorter into individual wells of a 384 well plate containing 1uL PBS + 0.1U/uL Superasin. Each plate was immediately snap frozen and stored at -80C until processing. All reagent delivery was performed with a high speed Biomek FXP liquid handling robot, and all reactions unless otherwise noted were kept at 4C. The following modifications were made to the SCRB-seq^95^ protocol for library generation. Primer sequences are consistent with the SCRB-seq publication. 384 well plates containing sorted nuclei were thawed at room temperature for 30 seconds before addition of 1uL of a 2uM primer mix per well (a common template switch oligo and a well-specific barcoded RT primer). Plates were incubated at 72C for 3 minutes, then immediately transferred to a 384 well metal block on ice. 3uL RT buffer (6.67U/ul Maxima H-(Thermo Fisher, EP0753), 1.67mM dNTP, 1.67x RT Buffer, 0.67U/uL Superasin, 1:5,000,000 ERCC spike-in) was added to each well. Plates were incubated at 42C for 90 minutes, followed by 10 cycles of 50C for 2 minutes and 42C for 2 minutes, with a final 70C inactivation incubation for 10 minutes. Plates were immediately transferred to ice, and 7uL of PCR mix (0.35uM SingV6 common forward and reverse primer, 0.033U/uL KAPA HiFi DNA Polymerase (KAPA, 07958846001), 1.67x HiFi Fidelity Buffer, 0.5mM dNTP) was added per well. PCR amplification was performed at 98C for 3 minutes, 18 cycles of [98C for 15 seconds, 67C for 30 seconds, 72C for 6 minutes], followed by a final 5 minute 72C elongation step and 4C hold. The number of PCR cycles had been optimized for this tissue using test plates. The 384 in-well reactions were pooled, purified with 0.8x AMPure XP (Beckman Coulter, A63881) according to manufacturer’s recommendation, and eluted in 50uL ultrapure water. 1ul of purified cDNA was used as input for Illumina Nextera XT tagmentation according to manufacturer’s recommendation, with a unique N7xx index for plate identification and a common i5 PCR primer (P5NEXT) for 5’ end cDNA-specific amplification. PCR cycles for library amplification were determined per library, and ranged from 12-18 cycles. Libraries were purified with one round of 0.8x ampure followed by 1 round of 0.65x ampure. Library structure and concentration for each plate was determined on a Bioanalyzer, and libraries from all plates were multiplexed for sequencing on a NextSeq 550 with the High Output 75 kit (17 cycles Read1, 8 cycles Index1, 58 cycles Read2).

#### STAB-seq Transposome Complex Assembly

40uM of a single ME-A_Calling Card oligo was annealed with 40uM ME-R oligo (5’Phosph-CTGTCTCTTATACACATCT) in 1x reassociation buffer (10mM Tris pH8.0, 50mM NaCl, 1mM EDTA), cooling from 98C to 4C at 0.1C/second. 40uM of a single ME-B_Calling Card oligo was annealed with 40uM ME-R oligo in the same way. The components for the transposome complex (TC) were pipet mixed on ice using 2.5ul of each annealed oligo, 7.5ul of 1.5x TC buffer (75mM HEPES-KOH pH7.2, 150mM NaCl, 1.5mM DTT, 0.15% Triton X-100, 15% glycerol), and 7.5ul of 0.3mg/mL pA-Tn5 (Active Motif, Cat#30721001). The TC was incubated at room temperature for 50mins, and then diluted to 1uM by adding 11ul of storage buffer (50% glycerol in 1x TC buffer). This 1uM TC can be stored for at least one month at -20C without loss of activity.

#### STAB-seq Assay

The following modifications were made to the single nuclei Cut&Tag^96^ protocol for antibody-directed nuclei tagmentation. Nuclei counting was performed on a hemocytometer with 405 wavelength by diluting the sample 1:2 with PBS + 0.1% BSA + 2ug/mL DAPI; DAPI was excluded from all incubation buffers. Nuclei were resuspended in Wash Buffer with EDTA (20mM HEPES pH7.5, 150mM NaCl, 0.5mM spermidine, 1x Roche EDTA-free protease inhibitor, 0.01% NP-40, 2mM EDTA), lightly fixed with 0.1% PFA for 2 minutes at room temperature, quenched by addition of glycine to a final concentration of 75mM, and pelleted at 500g for 5 minutes in a swinging bucket centrifuge. The following steps were performed in a 200uL volume, unless otherwise noted. Nuclei were washed once in 1mL Wash Buffer with EDTA, and resuspended in 400uL Wash Buffer with EDTA. 100k nuclei were used as input into STAB-seq. Nuclei were incubated overnight rotating at 4C with 1:50 primary antibody (H3K4me1 [Abcam, ab8895], H3K4me2 [Millipore Sigma, 07-030], H3K27ac [Active Motif, 39133], H3K27me3 [Cell Signaling Technology, 9733]). Nuclei were washed three times in Wash Buffer (500g for 5 minutes), then incubated 1 hour rotating at room temperature with Guinea Pig anti-Rabbit secondary antibody (Antibodies-Online, ABIN101961). Nuclei were washed 3 times in Wash Buffer (500g for 5 minutes), then resuspended in High Salt Wash Buffer (20mM HEPES pH7.5, 300mM NaCl, 0.5mM spermidine, 1x Roche EDTA-free protease inhibitor tablets, 0.01% NP-40) with 1:50 custom calling card-loaded pA-Tn5 TC (20nM) to uniquely label histone modifications, and incubated for 1 hour rotating at room temperature. Nuclei were washed three times in High Salt Wash Buffer (300g for 3 minutes), then resuspended in 100uL Tagmentation Buffer (High Salt Wash Buffer + 10mM MgCl2) and incubated shaking at 37C for 1 hour. Nuclei were pelleted (300g for 3 minutes), washed once with High Salt Wash Buffer (300g for 3 minutes), and resuspended to a final concentration of 3,640 nuclei/uL in PBS + 0.1% BSA. 60,000 nuclei were subjected to a second round of tagmentation with unbarcoded Tn5 using the BioRad SureCell ATAC-seq Library Prep Kit (BioRad, 17004620) according to manufacturer’s recommendation. After the second tagmentation, nuclei were pelleted (500g for 5 minutes), and half the supernatant was removed to concentrate nuclei to 2,400 nuclei/uL. Concentrated monodispursed nuclei were used as input into the ddSeq droplet generator, and custom 5uM STAB-N7xx indexing primers were used in place of the provided N7xx indexes. Manufacturer’s recommendations were followed for ddSeq droplet encapsulation, in-droplet indexing PCR (8 cycles), and amplified DNA purification. 2.5uL of the amplified product was input into qPCR to determine the appropriate number of second round PCR cycles. Libraries were amplified to log-linear phase (7-9 cycles) with the SureCell ATAC-seq Library Prep Kit reagents, purified with two rounds of 1x ampure, and run on a Bioanalyzer for quantification and structure assessment. Libraries were multiplexed for sequencing on a NextSeq 550 using High Output 300 reagents at 150x150x8 cycles, with custom Read 1 (BioRad SureCell ATAC-seq Library Prep Kit), and custom Read 2 and Index 1 sequencing primers.

#### Custom In-House Primer Sequences

ME-A_Calling Card 1

TCGTCGGCAGCGTCGCTAGACTAGATGTGTATAAGAGACAG

ME-A_Calling Card 2

TCGTCGGCAGCGTCTCGCTATCAGATGTGTATAAGAGACAG

ME-A_Calling Card 3

TCGTCGGCAGCGTCCTAGCTCAAGATGTGTATAAGAGACAG

ME-A_Calling Card 4

TCGTCGGCAGCGTCCAGCATACAGATGTGTATAAGAGACAG

ME-B_Calling Card 1

GTCTCGTGGGCTCGGTCGATCTCAGATGTGTATAAGAGACAG

ME-B_Calling Card 2

GTCTCGTGGGCTCGGGCTACACAAGATGTGTATAAGAGACAG

ME-B_Calling Card 3

GTCTCGTGGGCTCGGTATCAGCGAGATGTGTATAAGAGACAG

ME-B_Calling Card 4

GTCTCGTGGGCTCGGCTCGCAACAGATGTGTATAAGAGACAG

STAB-N701 Indexing Primer

CAAGCAGAAGACGGCATACGAGATTCGCCTTAGTTCAGACGTGTGTCTCGTGGGCTCGG

STAB-N702 Indexing Primer

CAAGCAGAAGACGGCATACGAGATCTAGTACGGTTCAGACGTGTGTCTCGTGGGCTCGG

STAB-N703 Indexing Primer

CAAGCAGAAGACGGCATACGAGATTTCTGCCTGTTCAGACGTGTGTCTCGTGGGCTCGG

STAB-N704 Indexing Primer

CAAGCAGAAGACGGCATACGAGATGCTCAGGAGTTCAGACGTGTGTCTCGTGGGCTCGG

STAB-N705 Indexing Primer

CAAGCAGAAGACGGCATACGAGATAGGAGTCCGTTCAGACGTGTGTCTCGTGGGCTCGG

STAB-N706 Indexing Primer

CAAGCAGAAGACGGCATACGAGATCATGCCTAGTTCAGACGTGTGTCTCGTGGGCTCGG

STAB-N707 Indexing Primer

CAAGCAGAAGACGGCATACGAGATGTAGAGAGGTTCAGACGTGTGTCTCGTGGGCTCGG

STAB-N708 Indexing Primer

CAAGCAGAAGACGGCATACGAGATCCTCTCTGGTTCAGACGTGTGTCTCGTGGGCTCGG

STAB-customREAD2 Sequencing Primer

GTTCAGACGTGTGTCTCGTGGGCTCGG

STAB-customIndex1 Sequencing Primer

CCGAGCCCACGAGACACACGTCTGAAC

#### *in situ* Sequencing by STARmap

*in situ* sequencing was performed following a modified version of the STARmap protocol. Briefly, tissue was sectioned at 16um onto poly-L-lysine treated coverslips, post-fixed, permeabilized, and hybridized with SNAIL primer and padlock probes overnight at 40° C. The following day, the probes were ligated using T4 ligase at room temperature and amplified using Phi29 polymerase at 30° C, both for two hours. Tissue was treated with BS(PEG)9 to facilitate cross-linking and anchored to a hydrogel matrix. Tissue was cleared of remaining proteins using Proteinase K for up to 24 hours at 37° C. Six rounds of sequencing and imaging were performed using a home-built fluidics setup based on previously described platform^97^, followed by detection of the polyA SNAIL primer and padlock probe and DAPI for cell segmentation.

#### Gene Selection and Probe Design

146 genes intersecting with cell type specific markers identified through snRNA-seq were selected for *in situ* profiling. A minimum of four unique genes for pan-neuronal, excitatory neuron, inhibitory neuron, motor neuron, pan-astrocyte, grey matter astrocyte, white matter astrocyte, astrocyte subpopulations 1-5, proliferative cells, pan-microglial, microglia subpopulations 1-4, pan-OPC, pan-oligo, and oligo subpopulations 1-2 were selected. Probes for STARmap were designed utilizing the PaintSHOP^98^ command line workflow, with modifications to account for the length of STARmap probes vs. MERFISH probes. Briefly, we specified a desired probe length of 42-50 nucleotides, 10% formamide concentration, hybridization temperature of 40° C (as input to NUPACK^99^ for structural analysis) and kmer length of 21 nucleotides (as input to Jellyfish^100^ to check for kmers). The output probes were split into their SNAIL primer and padlock constituents with a 2-3 nucleotide separation between them. Primer and padlock halves were separated such that their melting temperatures differed by no more than 2° C. These primer and padlock candidates were appended with the corresponding common sequence (including a unique barcode for each gene as part of the padlock probe) and run through PBLAT^101^ to eliminate probes mapping <17 nt to the coding region of another target. Of the primer/padlock pairs that remained, we manually mapped them via BLAT^102^ to ensure adequate separation between probe candidates (at least 100 nt between primer/padlock pairs) and to ensure that the constant regions of the primer or padlock probe did not map to the transcript of interest. If it was not possible to design four probes in an exon, due to transcript length, we supplemented the possible exonic probes with probes that met all criteria in the UTR. Separately, we designed a primer and padlock pair to map to the polyA tail of mRNAs to serve as a cell boundary marker. We utilized a separate common sequence backbone for the padlock probe to ensure no cross-talk occurred between this polyA probe and detection of other desired transcripts. This probe was detected after sequencing, using a universal detection probe complementary to this alternate padlock backbone attached to ATTO647 dye for visualization.

#### Coverslip Treatment

Cover slips were cleaned in an ultrasonic water bath by immersion in 2% RBS-35 (Thermo 27950) followed by 100% EtOH, washing three times with Milli-Q water in between, then allowed to dry in a 90° C oven. Cover slips were silanized by treatment with 1% Bind-Silane (γ-methacryloxypropyltrimethoxysilane; Cytivia 17133001) in acidic ethanol solution (95% EtOH supplemented with 5% glacial acetic acid) for one hour at room temperature, washed three times with 100% EtOH, and placed in a 90° C oven for at least 30 minutes to dehydrate the silane layer. Cover slips were further functionalized by 0.01% Poly-L-Lysine (Sigma P8920) in 1X PBS for three hours, washed three times with Dnase/Rnase-free water, and allowed to dry before use. Functionalized cover slips could be stored in a dessicated chamber for several days before use.

#### Tissue Preparation and Library Creation

*in situ* sequencing was performed using a modified version of the STARmap protocol^81^. Tissue sections were collected at 16um on a cryostat onto 40 mm round coverslips (Bioptechs) and post-fixed with 10% neutral buffered formalin (Sigma HT5011) at room temperature for 15 minutes, then permeabilized using 0.25% Triton X-100 for 10 minutes, followed by 0.1% pepsin in 0.1 N HCl for one minute. Three washes with 1X PBS were performed between each step. The tissue was dehydrated in an EtOH series; 50%, 70%, and 100% twice for 5 minutes each. The tissue was then allowed to fully dry on the cover slip before further processing. Tissue was re-hydrated in PBSTR, consisting of 1X PBS + 0.1% Tween-20 (Sigma 655204-100ML) + SUPERaseIn RNase inhibitor (Invitrogen AM2696) for 5 minutes, and then blocked using hybridization buffer without probes for 30 minutes at 40° C. Hybridization buffer consisted of 2X SSC, 10% formamide (Invitrogen AM9342), 20 mM ribonucleoside-vanadyl complex (RVC; NEB S1402S), 0.1 mg/mL salmon sperm DNA (Invitrogen AM9680), and 100 nM of the appropriate SNAIL probes, including the polyA primer and padlock probe if desired. After blocking, tissue was hybridized in 100 uL of hybridization buffer plus probes in a humidified chamber at 40° C overnight with gentle shaking. After hybridization, samples were washed twice with PBSTR for 20 minutes followed by one wash with PBSTR + 4X SSC for 20 minutes. All washes were performed at 40° C. The sample was then briefly washed once more with PBSTR at room temperature. Probe ligation and rolling circle amplification were performed as describe in the STARmap protocol. Briefly, wash buffer was exchanged with ligation mix, consisting of 1X T4 ligase buffer, 1X BSA (Invitrogen AM2618), 0.2 U/uL SuperaseIn, and a 1:50 dilution of T4 DNA ligase (Thermo Fisher EL0012). Ligation was allowed to proceed for two hours at room temperature with gentle agitation. Samples were washed twice with PBSTR at room temperature, then placed in rolling circle amplification mix, consisting of 1X Phi 29 buffer, 250 uM dNTPs (Invitrogen 18427088), 1X BSA, 0.2 U/uL Superase-In, 20 uM 5-(3-aminoallyl)-dUTP (Invitrogen AM8439), and a 1:50 dilution of Phi 29 DNA polymerase (Thermo Fisher EP0094). Amplification was performed for two hours at 30° C with gentle agitation. Tissue was washed twice post-amplification with PBST (no Rnase inhibitor). The addition of 5-(3-aminoallyl)-dUTP to the rolling circle amplification mix introduced a functional amine; we then treated the tissue with BS(PEG)9, a crosslinking agent, to facilitate cross-linking between probes and the hydrogel formed in the next step. BS(PEG)9 was resuspended in anhydrous DMSO as per manufacturer’s recommendation, then diluted to 50 mM in PBST. Treatment proceeded at room temperature with gentle shaking for one hour, then crosslinking was halted by treatment with 1 M Tris-HCl, pH 8.0 for thirty minutes. Polymerization buffer, consisting of a final concentration of 4% 19:1 Acrylamide/Bis-Acrylamide (Bio-Rad 1610144) and 2X SSC, was degassed under vacuum for 15 minutes. Separately, a 20% (vol/vol) solution of N,N,N’,N’-Tetramethylethylenediamine (TEMED; Sigma, T9281) and a 20% (wt/vol) solution of Ammonium Persulfate (AP; Sigma A3678) were prepared and kept on ice until use. The sample was rinsed thoroughly with 2X SSC and then equilibrated in degassed polymerization buffer for 30 minutes at room temperature. Immediately before use, TEMED and AP were added to the polymerization mix at a final concentration of 0.05% (vol/vol) and 0.05% (wt/vol) respectively, and a 40-uL droplet was placed on a Gel Slick (Lonza, 50640) treated glass slide. The cover slip with tissue was gently inverted onto this droplet, avoiding the formation of air bubbles, to form a thin hydrogel. Polymerization was allowed to occur at room temperature in a humidified nitrogen chamber (reference for seqFISH+) until oxygen was purged from the chamber (approximately 10 minutes), then moved to 4° C for 30 minutes, followed by 37° C for 2.5 hours. After hydrogel formation, the cover slip was gently detached from the glass slide and washed three times in PBST for five minutes each wash. Tissue was then digested at 37° C for up to 24 hours in a digestion buffer consisting of 50 mM Tris-HCl pH 8, 1 mM EDTA, 0.5% Triton X-100, 500 mM NaCl, 1% SDS, 30 mM glycine, and a 1:100 dilution of Proteinase K (NEB P8107S), changing the digest once. Tissue was then washed thoroughly with 2X SSC before imaging.

#### Imaging for in situ Sequencing

Tissue was stained with 1 ug/uL DAPI in 1X PBS for 10 minutes, then loaded into a Bioptechs FCS2 flow cell for imaging. Reagent delivery was performed using a home-built fluidics system. Briefly, we used an IDEX valve controller (MXX778-605) to select the reagent for delivery, which was pumped using negative pressure by a peristaltic pump (Gilson MP3) at the outlet of the flow cell set to pump at approximately 500 uL/minute. For the first round of sequencing, tissue was washed by PBSTG (PBST supplemented with 30 mM glycine to quench any remaining stripping buffer) then incubated in sequencing mix for 3 hours. Sequencing mix consisted of 1X T4 buffer, 1X BSA, 10 uM of round 1 reading probe, 5 uM of the fluorescent detection oligos, and a 1:25 dilution of T4 ligase. The sample was then washed with a wash buffer consisting of 10% formamide and 2X SSC, followed by GLOX imaging buffer consisting of 10% (wt/vol) glucose, 10 mM Tris-HCl pH 8, 2X SSC, 2 mM Trolox (Sigma 238813), 50 uM Trolox Quinone (reference for making TQ), 0.5 mg/mL glucose oxidase (Sigma G2133-50KU), and a 1:1000 dilution of catalase (Sigma C3515). Imaging was performed on an Andor Dragonfly spinning disk confocal system (talk about all the specs of the scope) using a Nikon Ti2 stand equipped with the Perfect Focus System (PFS) to maintain positioning of the sample during fluidics flow. We first imaged the tissue by DAPI to select the region of interest and ensure the uniform flatness of the imaging area. We then proceeded to image all four spot channels plus DAPI. For each subsequent round, the sample was incubated in stripping buffer (80% formamide with 0.1% Triton X-100), PBSTG, sequencing mix (supplemented with the appropriate readout probe), wash buffer, and imaging buffer, then imaged the sample. Six rounds of sequencing were performed, with four readout channels per round. After the sixth round of sequencing, the tissue was re-stained with DAPI and stained with a universal detection probe coupled to ATTO 647 to detect poly-dT signal for cell boundary segmentation.

#### Image Processing

Vignetting correction of each FOV was performed utilizing a non-parametric method as previously described.^103^ Since the non-uniformity of field illumination is relatively consistent across all fields of view in a single experiment, a small subset of FOVs across the imaging plane were selected to estimate vignetting correction parameters, and these parameters were averaged and applied to the entire data set. To determine vignetting correction parameters, each relevant FOV was downsampled by 4 across the x and y dimension to speed up processing, and a sigma value of 10,000 was used over 10 iterations. These calculations were performed separately across each imaging channel.

#### DAPI Segmentation

Cells were segmented by DAPI utilizing a multi-Otsu threshold and watershed transform provided by the scikit-image library. After thresholding, we performed a distance transform on the binarized images and looked for the peak local maximum of the distances within a certain minimum distance. Markers were generated from these local max labels. A watershed transform was then performed to segment each cell. Anything smaller than the expected pixel area for a single cell was thrown out to eliminate false positive cell detection. “Cells” detected that encompassed a pixel area far greater than expected for a single cell were flagged for closer examination, as they most likely were actually composed of more than one cell (under-segmented). We found the multi-Otsu thresholding approach helped to mediate between over- and under-segmentation of closely packed cells without adding excess complexity to the analysis pipeline. Re-thresholding under-segmented cells almost always yielded more accurate segmentation results.

#### oligo(dT) Segmentation

Cells boundaries were determined by segmentation of the polyA SNAIL probe signal. Images were first Gaussian blurred to smooth the spot puncta, then the same method as for DAPI segmentation was applied.

#### Stitching

Relative FOV positions and approximate locations across the imaged field were provided in an XML file generated by Fusion imaging software, and we utilized this information to provide a basis for FOV stitching. Stitching was performed on DAPI images that were vignetting-corrected, smoothed and downsampled by 4 in all spatial dimensions for memory conservation [to allow the entire dataset to be loaded into memory]. Although we knew the approximate location of each FOV, natural movement of the microscope stage during imaging introduced a small, few-pixel shift, so registration between overlapping regions of each FOV was necessary for smooth stitching. Registration was performed in the order of imaging based on that FOV’s overlap with the previous FOV. The estimated shifts between FOVs were also spot-checked by ensuring they were consistent with shifts between other neighboring FOVs. For example, if FOV1 and FOV2 overlap in all x pixels and 20% of y pixels, we can check that the determined shift is correct by registering the overlap between FOV1 and FOV2 over all y pixels and 20% of x pixels with the FOVs directly next to them. Segmented cells were stitched using these estimated parameters, and re-labeled to avoid duplication of cell labels across the entire stitched image. Detected spots were translated to their appropriate coordinates by FOV and shifted based on the parameters generated above. To avoid minor registration discrepancies and duplication of spots in overlapping regions as we built the stitched spots array, we only retained spots from the subsequent FOV in the overlap. A cell by gene counts matrix was then created by taking the union between spot locations and segmented cells.

### snRNA-seq and snATAC-seq Analysis

#### Pre-processing of 10X Multiome data

The 10X Multiome sequencing data for freshly resected thoracic and lumbar tissues were processed using the cellranger-arc pipeline. Specifically, raw base call (BCL) files were demultiplexed to generate fastq files. The “count” mode of cellranger-arc was then applied on each library. It should be noticed that by default “cellranger-arc count” considered the intronic reads when analyzing gene expression, which fitted our single-nuclei sequencing data. The outputs of “cellranger-arc count” in thoracic and lumbar were aggregated separately by “cellranger-arc aggregate”, which yielded two tissue-specific peak lists. We merged the peaks in the two lists and ran “cellranger-arc aggregate” again on all libraries in terms of the merged peaks. The output was a cell-by-gene count matrix for RNA, a cell-by-peak count matrix and a fragment file for ATAC. Cells from all libraries shared the same genes and peaks.

#### Pre-processing of Donor 1 snRNA-seq

Two FASTQ files were generated for each of the SCRB-seq plate, namely the R1 and R2 files. Reads in R1 file were 16-bp in length. The six nucleotides at the 5’-end of each read were cellular barcodes, and the following ten nucleotides were molecular barcodes. Reads in R2 file were the nucleotide sequences of RNA molecules. We aligned the reads in R2 file to the reference hg38 using STAR with parameters “--outSAMtype BAM SortedByCoordinate --outSAMunmapped Within –outSAMattributes Standard”. The cellular and molecular barcodes were appended to the alignment record of each read in the BAM file as the CB and UB tags, which made the BAM file capable to be the input of Velocyto. We ran Velocyto on the BAM file to generate cell-by-count matrix. The 384 SCRB-seq barcodes were input to Velocyto as the parameter “-b”, and the GTF file of human GENCODE v34 was input as the annotation file. The multiple mapped reads were automatically filtered out by Velocyto and did not add uncertainty to matrix generation. The output of Velocyto contained three matrices for each plate: spliced, unspliced and ambiguous. We took the sum of the three matrices as the cell-by-count matrix, while the spliced and unspliced matrices were also kept for the velocity analysis. In order to remove pseudo genes, we only kept the genes that intersected with RefSeq genes.

#### Pre-processing of Donor 1 snATAC-seq

We applied the Bio-Rad ATAC-seq analysis toolkit on the Bio-Rad single-nucleus ATAC-seq data. Specifically, we ran the FASTQ quality control, FASTQ debarcoding, alignment, alignment QC, bead filtration and bead deconvolution steps independently on every index. In the bead filtration step, we reviewed the curves of reads per barcode in every index. The knee points needed to be manually corrected in several indexes to make the number of retained cells close to expectation. After these steps, a demultiplexed BAM file with cell source information was generated for each index. In order to obtain a uniform peak list, we merged the BAM files from all indexes in all segments and called peaks using “macs2 callpeak” with parameters “-n all -f BAM --nomodel --keep-dup all --extsize 200 --shift - 100”. The cell-by-peak matrices were then generated by ChromVAR. We also converted the demultiplexed BAM files to fragment files using Sinto.

#### Batch Effect Correction

In order to achieve more robust and accurate cell type and subtype identification, we merged all 10X Multiome RNA data from freshly resected thoracic and lumbar tissues. We managed to create a non-negative cell-by-gene matrix that was free of batch effect using Liger. After running Liger until the OptimizeALS and QuantileAlignSNF steps, we obtained a cell loading matrix (H.norm) and a gene loading matrix (W) were created. According to the design of Liger, batch effect was corrected in QuantileAlignSNF step so that no batch effect existed in the cell loading matrix H.norm. We took the product of the two loading matrices H.norm and W to recover a non-negative cell-by-gene matrix for further analyses. In a similar way, we merged all the SCRB RNA-seq data from post-mortem cervical, thoracic and lumbar, and created a cell-by-gene matrix without batch effect for post-mortem tissues. For ATAC cells from freshly resected and post-mortem tissues, we calculated gene activity matrices using Signac and created batch effect corrected gene activity matrices using the same approach.

#### Clustering for Donors 2-6

We applied Scanpy to cluster the 10X Multiome RNA data from freshly resected tissues. Taking the abovementioned batch corrected expression profile as input, we identified the highly variable genes using the function scanpy.pp.high_variable_genes. Principal components were then calculated on the expression of the highly variable genes. We selected 30 principal components because they could explain more than 90% of the variance. Then we generated neighbor graph using scanpy.pp.neighbors with n_neighbors = 50, and performed leiden clustering under different resolutions (0.01, 0.02, 0.05, and from 0.1 to 1.5 by taking 0.1 as the increment step). Under each resolution, we calculated Silhouette score using the principal component matrix on top of the resulting clusters. We adopted the clustering results given by resolution = 0.1, which yielded 13 clusters and corresponded to maximal Silhouette score. The clusters were annotated by reviewing the expression or gene activity scores of general cell type markers in spinal cord. We further performed sub-clustering on each cell type to identify different subtypes. Since the difference between subtypes was not as distinct as the difference between major cell types, rigorous approaches should be carried out to ensure clustering stability. We perform sub-clustering following the paradigm proposed by Scclusteval, in which the clustering stability can be evaluated through sub-sampling. Below are the detailed steps:

1. Take out the expression profiles of the cells in a certain cell type, identify highly variable genes, calculate principal components and select the top principal components that explained at least 90% of total variance and calculated neighbor graph using n_neighbors = 50.
2. Perform leiden clustering under different resolutions (0.01, 0.02, 0.05, and from 0.1 to 1.5 by taking 0.1 as the increment step). Given a resolution *R*, we denote the set of cells in every cluster as *C*_*R*,*i*_, 1 ≤ *i* ≤ *K*_*R*_, where *K*_*R*_ is the number of clusters under resolution *R*. In order to ensure precise subpopulation detection, we aim at selecting a resolution that results in as many clusters while keeping stability. We evaluate the stability through subsampling in the next steps.
3. Randomly select 90% cells from the cell type, repeat step 1 and 2 on the subsampled cells. For each resolution *R*, denote the set of cells in every cluster as 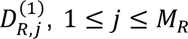, where *M*_*R*_ is the number of clusters on the subsampled cells. The superscript (1) in the notation 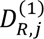 represents the first round of subsampling.
4. For each resolution *R*, calculate the maximal Jaccard index between every original cluster and the clusters on subsampled cells, which is defined as

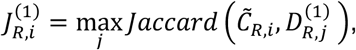

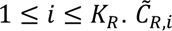 are the cells in *C*_*R*,*i*_ that appear in the subsampled set.
5. Repeat step 3 and 4 for *N* times. We set *N* as 100 in our implementation.
6. For each resolution *R*, calculate the stability score as

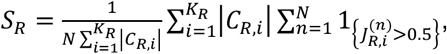

which is namely the proportion of maximal Jaccard index greater than 0.5 weighted by cluster size.

|*C*_*R*,*i*_| is the number of cells in *C*_*R*,*i*_. 1_{·}_ is the indicator function.

Our observation is that *S*_*R*_ has a general decreasing trend in terms of *R*. We select the 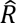 corresponding to the elbow point of *S*_*R*_, and the related clusters 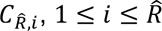 are taken as the subtypes.

#### Cluster Annotation for Donor 1

Given the obtained subtype labels for cells from freshly resected tissues, we inferred the subtypes for RNA cells from the post-mortem tissues using the optimal transportation plan between RNA-seq data. For each cell in the SCRB RNA-seq data, we went through the values in the optimal transportation plan matrix between that cell and every cell in the 10X Multiome RNA data. We selected the cells in 10X Multiome RNA data that corresponded to the top 20 values, and the most frequent subtype associated with the selected cells was taken as the subtype inference. In a similar way, we inferred subtypes for the ATAC cells from post-mortem tissues using the optimal transportation plan between the SCRB RNA-seq data and Biorad ATAC-seq data.

#### Canonical Correlation Analysis

In this section, we let *X* ∈ ℝ^*n*×*p*^ and *Y* ∈ ℝ^*m*×*p*^ denote the two data sets obtained after data preprocessing [cf Anqi’s part], where *n* and *m* are the numbers of cells in the first and second data sets respectively, and *p* is the number of genes. We first embed *X* and *Y* in a common, low-dimensional common space *Z* using Canonical Correlation Analysis (CCA). The objective of CCA is to find a linear projection of the data such that the correlation matrix *C* = *corr*(*X*, *Y*) is as large as possible, where correlation is defined between two (empirical) random variables *x*, *y* ∈ ℝ^*p*^ as

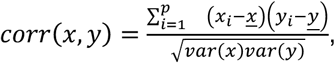

and 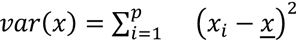. In particular, CCA aims at finding projection matrices *A* ∈ ℝ^*n*×*d*^ and *B* ∈ ℝ^*m*×*d*^, for some low dimension *d* ∈ ℕ^∗^, such that *corr*(*A*^*T*^*X*, *B*^*T*^*Y*) is maximized. In practice, we compute the correlation matrix *C* ∈ ℝ^*n*×*m*^ between our two data sets, and we then compute a singular value decomposition of *C* : *C* = *A*Z*B*^*T*^. The left and right singular vectors *A* ∈ ℝ^*n*×*d*^ and *B* ∈ ℝ^*m*×*d*^ of this SVD provide the two embeddings that maximize the correlation. In our analysis, we set *d* as the dimension for which the explained variance achieves 99 % of the total variance.

### Optimal Transport

Once CCA has been used to embed *X* and *Y* in a common embedding space, we use Optimal Transport (OT) to align the cells. OT is a very common tool of applied mathematics that allows to compare discrete probability measures by finding an alignment, or correspondence, between the support of the measures. More formally, given a space *X* endowed with a cost function *c*: *X* × *X* → ℝ_+_, and two discrete measures *μ* and *v* on *X*, namely measures that can be written as positive combinations of Dirac measures, 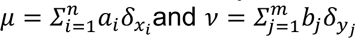 with weight vectors 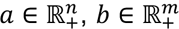 satisfying *Σ*_*i*_*a*_*i*_ = *Σ*_j_*b*_j_ (i.e., the measures have same total masses) and all *x*_*i*_, *y*_j_ ∈ *X*, the *n* × *m* cost matrix *C* = (*c*(*x*_*i*_, *y*_j_))_*ij*_ and the set of candidate transportation matrices, defined as

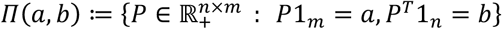

define a so-called optimal transport problem. The optimal transport plan *P*^∗^can be computed using the following linear program:

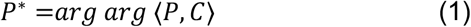

where ⟨⋅,⋅⟩ is the Frobenius dot product, i.e., 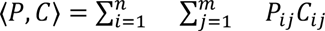. Unfortunately, it is well-known that solving the optimal transport problem is intractable when data sets are large. In particular, our single–cell data are too large to compute the optimal solution *P*^∗^ exactly. A common and very efficient workaround^104^ is to consider an entropic regularization of the optimal transport problem, namely:

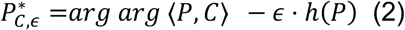

where *∈* > 0 and the negentropy ℎ is defined as *h*(*P*) ≔ −*Σ*_*ij*_*P*_*ij*_(*log log* (*P*_*ij*_ – 1). Since the negentropy is strongly convex, the regularized optimal transport problem admits a unique solution, and can be computed efficiently. Indeed, it is known that *P*^∗^ takes the following form:

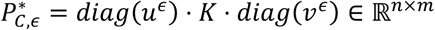

where *K* is computed by exponentiating each term of *C* with 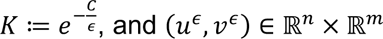 can be computed as the fixed points of the so-called Sinkhorn map: 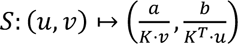.

Note that these fixed points are the limits of any iterative sequence (*u*_*t*+1_, *v*_*t*+1_) = *S*(*u*_*t*_, *v*_*t*_), which immediately gives an algorithm to estimate *P_C,∈_*^*^, known as Sinkhorn iterations. The Sinkhorn divergence is defined as the transport cost of the optimal regularized plan, 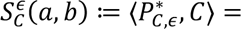 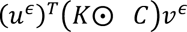 (where ⊙ denotes the term-wise multiplication), and is known to converge 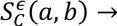 ⟨*P*^∗^, *C*⟩ as *∈* → 0, and more precisely *P*^∗^*_C,*∈*_* converges to the optimal transport plan solution of (1) with maximal entropy. Finally, OT can be generalized to unbalanced OT whenever (2) is augmented with two Kullback-Leibler terms:

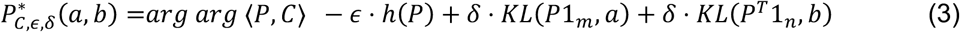

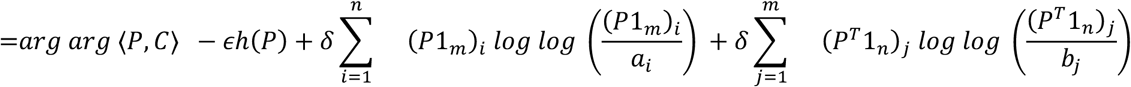

where *P* ranges now over the set 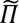 of the positive *n* × *m* matrices. Again, cost (3) can be solved using Sinkhorn iterations. In our analysis, we always use unbalanced OT between our data sets 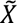 and 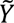 preprocessed with CCA, using the Euclidean pairwise distance matrix 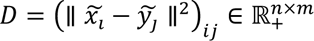 between 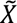 and 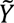 as the cost matrix *C* ≔ *D*, and uniform weight vectors 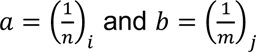. The entropic and marginal regularizations *ε* and *δ* are chosen in the list

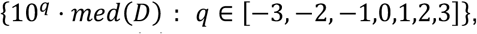

where *med*(*D*) is the median of D. More precisely, *ε* and *δ* are chosen as the smallest values in that list such that numerical errors are avoided. OT transportation plans are computed with the POT Python package. Once an optimal transportation plan *P*^∗^ ∈ ℝ^*n*×*m*^ has been computed, we use it to transfer information (such as, e.g., cell types). For a given cell 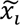, we aggregate the k-th largest values and their indices

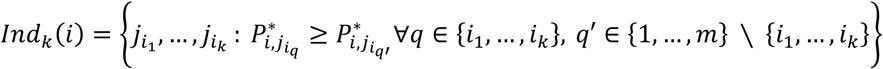

of row *i* in matrix *P*^∗^, and select the most frequent information associated to this subset. Moreover, in order to avoid selecting an arbitrary *k* for transferring information, we run this transfer for *k* ∈ {5*k*′ : *k*′ ∈ 2, … ,20}, and select again the most frequent information among all transferred ones (one for each value of *k*).

### GRN analysis

#### Inferelator summary

The Inferelator method derives Gene Regulatory Networks (GRNs), elucidating interactions between Transcription Factors (TFs) and genes, by integrating regulatory evidence and expression data^105^. Regulatory evidence comprises a prior binary matrix of regulatory links between genes and TFs, derived from evidence combining accessible chromosomal elements with TF. An intermediate computed state of the Inferelator is TF activity (TFA) estimates that can be used for cell specific analysis of regulatory effects of TFs. For the expression data, each cell is normalized to a count of 10^4^. Genes expressed in >100 nuclei are retained. Subsequently, expression values undergo log transformation with the addition of a pseudo count using Scanpy’s log1p function^106^. For the analysis, samples from Donors 2 and 3 are excluded due to an insufficient number of sampled cells for both Thoracic and Lumbar regions across all relevant cell types.

#### Lumbar Thoracic TFA

When constructing the regulatory evidence prior, each gene is mapped to a TF using the Inferelator-prior pipeline, TF motifs, and peaks from the accessome derived from snATAC-seq. A gene is associated with a TF if any peak has a matching TF motif. The prior is then filtered to enrich well-matched motifs, retaining at most 5% of links. In this instance, 4.75% of the total possible prior associations are retained using a motif match score threshold of 40. GRN inference with the Inferelator using the multi-task amusr workflow, splitting the data into two tasks: thoracic and lumbar. A unique prior for each segment is constructed by including evidence only from accessible peaks in that segment, resulting in two distinct priors. For thoracic and lumbar segments, there are 25,304 and 25,270 genes respectively, with 273 TFs common relevant to both segments. To ensure robust network estimates, the data is bootstrapped 10 times. Transcription Factor Activity (TFA) estimates are computed for each TF and cell using task-specific priors according to the model:

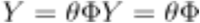

where *Y* ∈ *R^S,n^* is the gene expression matrix with n genes and S samples, Φ ∈ *R^m,k^* is the TFA with k TFs, and Θ ∈ *R^k,n^* is the GRN. Φ is unknown and to deconvolve and find TFA estimates we impose a task specific prior *P_C_* with elements ∈ 0, 1 and use that to solve for an estimate of TFA;

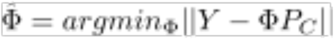

#### Differently Active TF Estimates

Using Scanpy’s rank_gene_groups function, a t-test is performed for each cell type, comparing activity between the Thoracic and Lumbar segments. A Benjamini-Hochberg adjusted p-value of 10^2^ is employed for all differentially active TF estimations unless otherwise specified.

### STAB-seq Data Analysis

#### STAB-seq Data Pre-Processing

We processed the STAB-seq data using Bio-Rad ATAC-seq analysis toolkit, while additional steps were necessary to parse the calling cards and remove crosstalk. Below are the major steps:

- Parsing calling card: To ensure successful running of Bio-Rad ATAC-seq analysis toolkit, we removed calling cards that were inserted in any sequencing reads. However, we assigned each read pairs a tag to record the presence of calling cards, which can be “dual” (both R1 and R2 mates had calling cards), “singleA” (only R1 mate had calling card), “singleB” (only R2 mate had calling card) and “unbarcode” (none of R1 or R2 mate had calling card).
- Running Bio-Rad analysis pipeline: We input the fastq files with calling cards being removed to Bio-Rad ATAC-seq analysis toolkit and ran through the cell filtration step. Three files were used in following steps: (1) BAM file generated by the alignment step; (2) BAM file generated by cell filter step; (3) TXT table generated by the deconvolution step that recorded the bead barcode and cell barcode translation. The major difference between the two BAM files was that reads in the latter were deduplicated.
- Decrosstalk: The aim of this step was to filter out the crosstalk which occurred in PCR amplification that a calling card was wrongly inserted to a molecule where it should not appear. We started with adding a combinatorial tag to each read in the BAM file generated by the alignment step. The combinatorial tag was in the format of “Cell_barcode+Calling_card_tag”, where the cell barcode was obtained by looking up the bead and cell barcode translation table and the calling card tag was saved in the calling card parsing step. Next, we applied Sinto on the BAM file to create fragment file. The output of Sinto included coordinates of each fragment and the most likely combinatorial tag associated with it. If a fragment had multiple combinatorial tags, Sinto assigned the one supported by the maximal number of read pairs, thus reducing the false positive that a fragment from the second tagmentation being identified as a modification or binding region. Finally, we used the fragment file to further filter the BAM file generated by the cell filter step of Bio-Rad pipeline. A read pair was kept only if the covered region was consistent with a region in the fragment file. The associated combinatorial tag was assigned to the read pair to indicate the cell barcode and calling card tag. We split the filtered BAM file by the calling card tag and kept the two tracks associated with “dual” and “unbarcode” for subsequent analyses.

#### STAB-seq Annotation

We merged the “dual” and “unbarcode” BAM files of the 27 STAB-seq libraries and called peaks using MACS2 with parameters “—nomodel --nolambda --keep-dup all”. Peaks were extended by 150 bp at both ends and overlapped peaks were merged. Then we used ChromVAR to create count matrices for each “dual” and “unbarcode” BAM file. We filtered out the cells whose total “unbarcode” fragment counts were no larger than 100, and also filtered out the cells in H3K27ac, H3K4me1 and H3K27me3 libraries in which the percentage of “dual” fragments were smaller than 5% or greater than 80%. To generate a UMAP representation of STAB-seq data, we combined all the “unbarcode” matrices and ran Signac with Harmony batch effect correction. We did not use the first two LSIs when creating UMAP because they had high correlation with total fragment counts (absolute values of correlations greater than 0.5). To annotate the STAB-seq data by transferring the cell type labels from 10X Multiome ATAC-seq data through optimal transportation. Specifically, we merged the peak lists of STAB-seq and 10X Multiome ATAC-seq and recreated count matrices of the data from the two modalities so that they shared the common peak list. Then we used Signac to generate LSI representation of the recreated count matrices and Harmony was called to reduce batch effect. We created an optimal transportation plan between the two modalities using the resulting LSIs. For each STAB-seq cell, we found the 100 10X ATAC-seq cells whose optimal transportation values ranked at the top. The most frequent cell type among the 100 cells was taken as the cell type inference for the STAB-seq cell.

#### Peak Calling

Fragment files (pre-filtered for cells) generated by the Bio-Rad pipeline were reprocessed using ArchR (v1.0.3 dev branch)^107^. After manual inspection of quality control metrics, cells were further filtered for TSSEnrichment ≥3 and nFrags ≥10^2.75^ for ATAC, H3K27ac, H3K27me3, and H3K4me1 modalities. A TileMatrix was populated with insertion counts at 500bp non-overlapping windows with ArchR::addTileMatrix and gene activity scores (GAS) were calculated using the ArchR::addGeneScoreMat function. Latent semantic indexing (LSI) was used to reduce dimensionality of the TileMatrix using ArchR::addIterativeLSI. Uniform manifold approximation and projection (UMAP) was performed on the LSI reduced dimensions for visualization. Group-wise peak-calling was performed with MACS (v2.2.7.1)^108^ according to RNA major cell-type labels and reduced into a non-overlapping set, as previously described^109^, using ArchR::nonOverlappingGR.

#### Marker Discovery

To simplify analysis, we used group-wise, accessible peaks as the search space for marker discovery for all modalities. Cell-type specific marker peaks were detected using ArchR::getMarkerFeatures using a binomial test, correcting for bias with ReadsInPeaks with 2000 maxCells, using binarized insertion counts from the “PeakMatrix”. Significant peaks (FDR < 0.1, Log2FC > 1) were selected. Marker peaks were evaluated for each of the relevant contrasts: 1) one vs all other cell types, 2) Astrocytes vs. Microglia, 3) OPC vs. Oligodendrocytes, 4) OPC vs. Oligodendrocytes (branch 1) vs. Oligodendrocytes (branch 2). Motif activities were calculated using the ArchR::addDeviations function, which adopts previously applied methods for estimating per cell variation in accessibility at motif-containing peaks against a GC and total accessibility matched background peak sets using chromVAR::getBackgroundPeaks^110^. Deviation Z-scores represent motif activities and were evaluated for cell-specific change to identify marker transcription factors. For each motif, the mean difference in motif activity was further scaled across cell types for visualization.

#### Imputing Gene Expression in STAB-seq

Given the STAB-seq data lacks expression information, we used the optimal transport plan, aligning cell-to-cell chromatin accessibility profiles, to assign the nearest 10X Multiome cells to each STAB-seq cell. The aggregated expression counts for each STAB-seq cell was then defined as the weighted aggregate, by OT distance, of the 50 nearest 10X Multiome cells. In order to minimize the sparsity in the STAB-seq signal, we also aggregated counts from the 50 nearest STAB-seq cells in the STAB-seq ATAC LSI reduced dimension space for each cell-type and modality.

#### Assessing STAB-seq Intermodal Associations

To assess associations between STAB-seq modalities, pseudo-bulk insertion pileups were aggregated (bigWig format)–100bp tiles, 1000 max cells per group, 4 max counts per cell, normalized by “ReadsInTSS”–using the ArchR::getGroupBW function. *Spearman rho* was calculated for all pairwise pileup comparisons and used as a distance (*1 - rho*) for clustering (WardD2) to demonstrate expected relationships between modalities. To evaluate co-occupancy of H3K27ac, H3K27me3, and H3K4me1 at noncoding peaks, we first annotated peaks using ChIPseeker::annotatePeak (prioritizing exons, UTR’s, introns, downstream, promoters, then intergenic)^111^. Insertion signals were extracted from pseudo-bulk bigWigs at noncoding peaks (excluding promoters, exons, and first introns). The signal mean for each group and modality was then converted into quantiles and the density difference between groups was visualized in a ternary plot.

#### SCARlink Modeling

In order to model the *cis*-regulatory effects of proximal accessibility and histone occupancy at accessibility peaks on expression, we used the SCARlink algorithm^34^ (https://github.com/snehamitra/SCARlink). To apply this model to our specific study context, we made minor modifications to the code to take peak coordinates instead of tiles as input and to allow for both positive and negative coefficients, i.e. for modalities where regulatory effects are known to be bidirectional or unknown–H3K27me3, H3K4me1. We converted our aggregated counts matrices for RNA (imputed from OT mapping of 10X Multiome to STAB-seq) and each STAB-seq modality into the necessary HDF5 input format (coassay_matrix.h5) for SCARlink. Matrices associating peaks within 100kb of the top differentially expressed (resources/DE) and known marker (resources/cell type markers.csv) genes for each contrast and modality combination were processed. To build per gene models, SCARlink was run on NYGC’s on-premises high-performance compute cluster with the Slurm Workload Manager for each modality and contrast, scarlink --celltype <CELL_LABEL> --outdir <OUTDIR> --genome genes.gtf --proc <SLURM_ARRAY_TASK_ID> --nproc <NJOBS> --sparsity .9. Subsequently, models were assessed for gene-linked peaks, scarlink_tiles --celltype <CELL_LABEL> --outdir <OUTDIR>.

#### Modality Integration

SCARlink models were filtered to identify peaks with a significant effect and peaks with no noticeable effect on expression. To do so, we classified peaks using the following constraints:

1. Fraction of non-zero cells was greater than 0.1 (test_acc_sparsity > 0.1)
2. Spearman correlation of predicted vs. observed expression > 0.1
3. Peak is a marker in the matched cell-type/modality (by ArchR::getMarkerFeatures) with a fold change >1.5X
4. Non-zero regression coefficient

Peaks satisfying these four criteria were deemed significant. Peaks for which the model fit was acceptable (Spearman correlation > 0.1) but the magnitude of peak change was small (fold change<1.25X) for all tested peak-gene pairs were deemed not significantly changed. All peaks modeled with SCARlink were filtered to only those that were significant in at least one modality and cell-type or were deemed not meaningfully changed. Descriptive classifications were given to significant peaks based on the context and relationship with expression. Classifications were split on whether associated with a peak with accessibility correlated with expression (*Co-accessible*) or no meaningful change in accessibility (*Accessible-independent*). Further, integrated classifications were aligned with expression direction for each of the contrasts: 1) one vs. other, up- or down-regulated in the cell-type of interest; 2) Astrocyte vs. Microglia up-regulated; 3) OPC vs. Oligodendrocyte up-regulated. The following table describes the classifications used where “+” denotes a significant positive coefficient, “-” denotes a significant negative coefficient, blank cells are insignificant, and “*” permits any coefficient including nonsignificant.

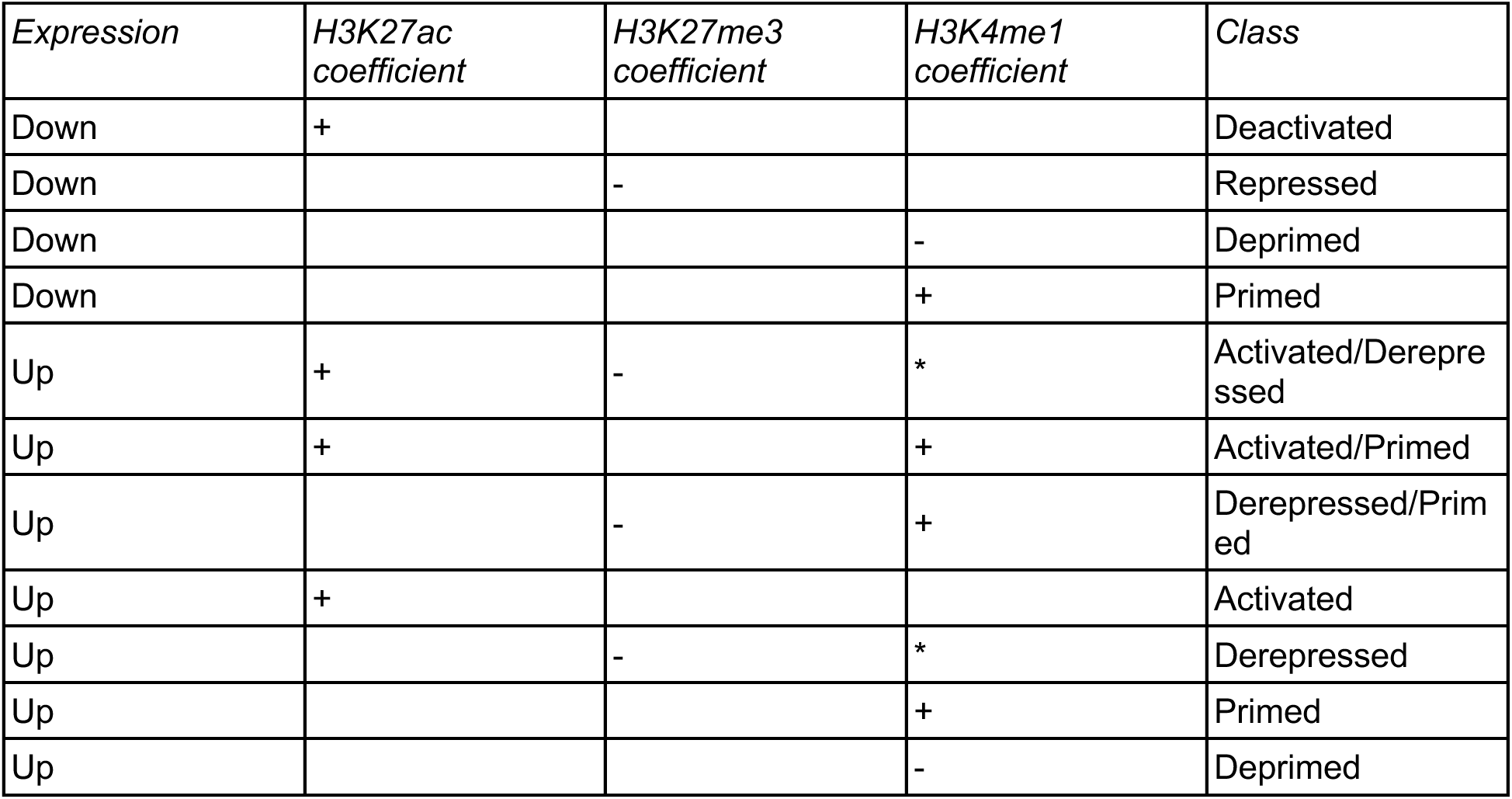

### scTDA Analysis

#### Detecting Patterns of Continuous Change

Focusing on the OPC and oligodendrocyte subpopulations, we wanted to extend our discrete clustering results to a continuous interpolation between various cell states. We reasoned that probabilistic versions of usual clustering algorithms may be able to uncover these gradients of change as they return values that reflect the strength of association of a point with a cluster on a continuous spectrum. An example is non-negative matrix factorization (NMF), which is a convex relaxation of the classical *k*-means algorithm. In general, matrix factorization methods have been used to great effect in analyzing single cell omics data, for example in identifying gene activity programs or performing batch integration.

However, the NMF algorithm, for example as implemented in scikit-learn, depends on a random seed that changes with each instance, which may hamper reproducibility and interpretability. In order to obtain robust factors from NMF, we used a bootstrap-like procedure called consensus NMF (cNMF) which combines multiple runs of the standard NMF algorithm on random subsets of the data to construct consensus factors.

Next, cNMF, like NMF, also requires a user-defined value of *k*, the number of factors used to decompose the matrix. As we will be using these factors to build topological representations and identify terminal cell states, we are somewhat limited in the number of factors we can use; we only need to find the strongest gradients in this step, which are generally very stable. Nonetheless, we generalized the cluster stability evaluation method, to handle probabilistic cluster assignments by replacing Jaccard similarity with Ruzicka similarity (), which is defined for two vectors *x* = (*x*_*i*_) and *y* = (*y*_*i*_) with non-negative real entries as

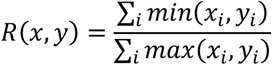

Note that if we restrict the vectors to have only binary values 0 or 1, this formula reduces to the Jaccard similarity. We used this metric to weigh the tradeoff between factor localization specificity and factor stability.

For both the postmortem tissue and fresh tissue datasets, we ran NMF with *k* ≤ 5 on the snRNA-seq inner product data for 100 trials, and constructed consensus factors using 20 of the trials, leaving the remaining 80 trials to assess factor stability. In subsequent analysis, we used the three factors from the cNMF results with *k* = 3 for the postmortem tissue dataset, and three stable factors from the cNMF results with *k* = 5 for the fresh tissue dataset.

#### scTDA Graph Representations

We refined the continuous patterns discovered using cNMF to create a continuous representation of OPCs and oligodendrocytes as a network. Many approaches for graph representations of single cell data have been proposed, ranging from simple *k*-nearest neighbor graphs and ɛ-neighborhood graphs to more complicated methods such as UMAP and PAGA. In this work, we decided to use Mapper graphs for their flexibility and their ability to handle arbitrary topologies in an unsupervised manner. In biology, Mapper graphs have previously been used to study cellular differentiation in single cell transcriptomic data of mouse embryonic stem cells, as well as to identify significant somatic mutations in cancer from bulk RNA-seq data.

We briefly describe the Mapper construction, which is based on the notion of partial clustering motivated from Reeb graphs and constructions in Morse theory. In addition to the input data *X*, Mapper uses a lens function *f*: *X* → *S* to determine the important topological features to emphasize and an overlapping covering 𝔘 of *S* to set the resolution scale. Given this information, we first pull back the covering along the lens function to obtain a data-aware covering *f*^−1^𝔘 of *X*. Next, for each subset *V* in the pullback covering *f*^−1^𝔘, we cluster the points in *V* using the metric in the original space *X*. Each cluster thus found becomes a node in the Mapper graph, and two nodes are joined by an edge if and only if their clusters overlap. In fact, we created a weighted version of the Mapper graph where the edge weights are determined by the size of the overlap. We used the software implementations KeplerMapper and NetworkX to generate and process Mapper graphs.

Depending on the filter function, the clustering algorithm, and other parameters, the Mapper construction can produce a plethora of different graph representations at different resolutions. We explain the inputs and parameters we used for the construction of Mapper graphs below.

- *Input data*: For each dataset, we used all the nonzero principal components obtained from the inner product data after filtering for highly variable genes using Scanpy. The Mapper construction implicitly performs its own dimensionality reduction so there is no need to further reduce the data beforehand up to moderately large dimensions of the ambient embedding space. We give the input data the structure of a metric space for clustering purposes using correlation distance.
- *Lens function*: For each dataset, we used the three cNMF factors found in “Robust discovery of patterns of continuous change in single cell sequencing data” as lens functions.
- *Clusterer*: We kept the default clustering algorithm in KeplerMapper, which is DBSCAN. We found DBSCAN to be a good choice since it is fast and also because it allows for the possibility of creating just one cluster if the points are sufficiently similar to each other as well as leaving outliers unclustered, which helps control the number of nodes in the resulting Mapper graph.
- *Clusterer parameters*: There are two main parameters for DBSCAN, the neighborhood size ɛ and the minimum points *minPts* in a neighborhood for a point to be considered a core point. A useful heuristic is to set *minPts* = *k* + 1, where *k* is equal to twice the intrinsic dimension of the data minus one, and to choose ɛ by locating an elbow in the *k*-distance plot. We generally followed these recommendations, but we also adjusted these values based on a stability analysis described below.
- *Cover parameters*: We covered the codomain of the lens function with overlapping axes-aligned rectangular boxes. More precisely, we used a product of regularly spaced intervals in each dimension as our open cover, which thus can be entirely described by two parameters: the number of bins *numBins* along each dimension and the overlap fraction *percOverlap* between adjacent bins. Here, our principal desideratum was to obtain a Mapper graph that is connected; a secondary priority was to resolve the expression space as finely as we could. The first criterion can be achieved using a small number of bins and a large overlap fraction, while the second leads to the opposite.

In order to finalize our choices of parameters for the Mapper algorithm and to assess the stability of the resulting graphs, we generated Mappers across a range of the parameters discussed above, and evaluated them based on connectivity, granularity, and topological consistency. To quantify the latter, we computed the correlation between the normalized internode graph distances between landmark nodes found using the procedure in “Data-driven identification of landmark nodes”, and picked parameters contained in a large region of the parameter space with correlation values *R* > 0.9. Ultimately, we chose ɛ = 0.3, *minPts* = 10, *numBins* = 15, and *percOverlap* = 0.55 for both the postmortem tissue data and the freshly resected tissue data, but the overall topologies of the Mapper graphs constructed with these choices are generally robust to perturbations of these parameters.

For visualization, we used the SFDP graph layout algorithm provided by Graphviz.

#### Multi-Branch Pseudotime Inference

A popular approach to inferring pseudotime from a graph representation is to use some version of a Markov process, also referred to as diffusion or a random walk process. The basic idea is that cells start in some node of the graph and transit along edges around the graph according to some probabilistic law. The aggregate motion of many cells gives rise to trajectories in the transcriptomic landscape that are parametrized by pseudotime.

However, this general description belies the many possible assumptions on the process needed to extract a pseudotemporal ordering, and different methods have been concocted to handle different cases depending on the availability of biological priors. One important distinction between the various methods is the topology that the method can handle, which range from simple linear trajectories all the way to disconnected graphs with cycles. For example, the Mapper algorithm makes no assumption on the topology, and so a priori can discover any type of trajectory. However, we make the assumption that the graphs we work with are connected, so that random walks on them are irreducible: any node can reach any other node via a path in the graph. The other piece of biological information we will use concerns the directionality of the trajectories. There is an inherent symmetry in pseudotimes constructed based on the similarity of transcriptomic profiles: reversing pseudotime yields another ordering that would explain the progression of transcriptomic changes just as well. Thus, there is a need for a method to break this symmetry; one common way this is done is by specifying root and terminal nodes. We describe one procedure for making these selections below.

To summarize, suppose we are given a connected graph representation of the expression landscape together with a set of root nodes and a set of terminal nodes. In fact, these nodes can be specified probabilistically instead, but to ease the exposition we restrict to the case where these root and terminal regions are localized at individual nodes. Using this information, we build a Markov process whose states are the nodes of the graph and the transition probabilities given by edge weights normalized to sum to one. For instance, recall that for the Mapper graphs the edge weights are given by the sizes of overlaps between clusters, which we now interpret as empirical measure of the number of cells that flow between the two nodes. Edges with small weights correspond to rare transitions, while edges with large weights correspond to frequent transitions. A variant of this could also take into account the size of the node as a proxy for the self-transition probability of that node, but we do not pursue this further here.

Now, we use the root and terminal nodes to modify the Markov process so that the root and terminal nodes are absorbing states, and compute the absorption probabilities of this process. These probabilities represent the time to absorption for a cell starting at a given node and ending at one of the absorbing nodes.

In more detail, let

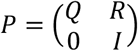

be the transition matrix for this absorbing Markov chain. We have closed-form expressions for the various properties of this process (Kemeny and Snell, 1976). For instance, the expected time to absorption (at any absorbing node) is *N*1, where *N* = (*I* − *Q*)^−1^ is the fundamental matrix of the Markov chain. Separating this out into absorption probabilities at each absorbing node, we let *B* = *NR*.

Then, the columns of the matrix *B* are the absorption probabilities starting at a non-absorbing node and ending in the absorbing node corresponding to the column.

Now, assume moreover that we have a unirooted process, i.e., there is only one root node. In this case, we can further simplify the topology of the system into a rooted tree with leaves corresponding to the terminal nodes. In this case, the relative absorption probabilities starting at a given node are an indicator for the “branch” to which the node belongs. Furthermore, the complement of the absorption probability at the root, or equivalently the sum of all the non-root absorption probabilities, is a measure of the global pseudotime distance from the root to the terminal states.

To increase the stability of and assess the robustness of these pseudotime results, we generated an ensemble of 1,000 Mapper graphs, each time using only 70% of the inner product gene expression data for OPCs and oligodendrocytes, and repeated the pseudotime inference procedure described above on the Mapper graph replicates. The node-level absorption probabilities were then transferred to individual cells and averaged over all the trials. The mean absorption probabilities were then normalized at the root, so that the average cell near the root has equal absorption probabilities at any of the terminal states; here the raw absorption probability of the “average root cell” was set to the median of the bottom 2% of cells with the lowest global pseudotimes. Other prior information about the eventual fate probabilities at the root can also be incorporated instead. Each cell was assigned to the branch (OPC, oligodendrocyte branch 1, or oligodendrocyte branch 2) for which its mean absorption probability was highest.

Finally, we grouped the cells within each branch and converted the mean absorption probabilities into quantiles – a monotonic transformation that preserves the ordering – which we interpret as a branch-specific pseudotime taking values between 0 and 1. For heatmap visualizations, we used *csaps* (github.com/espdev/csaps) with a smoothing parameter of 0.99 to create natural cubic smoothing splines for the gene expressions of the pseudotime-ordered cells in each branch.

#### Data-driven Identification of Landmark Nodes

We sought to automate the identification and selection of root and terminal nodes on a graph from the data. For terminal nodes, we observed that the values of the gradients found by cNMF are precisely maximized at the ends of each branch in the Mapper representation, and so we simply marked those nodes for which each of the factors from cNMF is highest as terminal. For the root node, we first computed the transcriptional entropy for each cell: if *p* = (*p_g_*) is the expression vector of highly variable genes for a cell, then the transcription entropy of the cell is

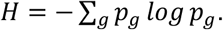

The node in the graph with the highest average transcriptional entropy was set as the root node.

### STARmap Analysis

#### High Depth Gene Expression Inference

We started with two rounds of quality control on the STARmap data. In the first filter, only cells with gene counts greater than 5 and smaller than 100 were kept. Then we calculated the median and standard deviation of gene counts across all the kept cells. In the second filter, we removed cells whose gene counts were outside the range of double standard deviation from the median. It should be noticed that these filters also removed 39 motor neurons which we manually identified. We added these motor neurons back and obtained 37,598 cells in total. After quality control, we applied the STARmap data analysis pipeline to perform clustering on the data. The number of principal components and nearest neighbors was set as 10 and 30, respectively. On the other hand, we performed non-negative matrix factorization on the count matrix using Liger and then took the product of cell loading and gene loading matrices, which created a smoothed expression profile. We built an optimal transportation between the smoothed STARmap data and 10X Multiome RNA-seq data. For each STARmap cell, we selected the top one hundred 10X RNA-seq cells indicated by optimal transportation. The most frequent cell type among the 100 cells was taken as the cell type inference for the STARmap cell. We also calculated the weighted sum of the log transformed TPM across the 100 cells and took it as the inferred expression for the STARmap cell. Due to the sparsity of motor neurons, we performed a precise optimal transportation between the 39 motor neurons in STARmap and 35 motor neurons in 10X RNA. For each motor neuron in STARmap, we took the best matched 10X RNA motor neuron as indicated by the optimal transportation for expression inference. It should be noticed that the cells in one of the STARmap cluster had obviously smaller read counts. Guided by the expression of marker genes, we did further filter on this cluster and only kept those cells that were inferred to by microglia and OPC by the optimal transportation. The total number of STARmap cells is 36,163 after this filter.

#### Community Detection

In this section, we explain how we get communities of cells in spatial data. We recall that spatial data is given as a cell by marker matrix *X* ∈ ℝ^*n*×*p*^, and a spatial coordinate matrix *C* ∈ ℝ^*n*×3^. Even though cell types can be inferred from marker gene expression in *X*, the (relatively) small number of markers does not allow for precise assessment of subgroups, and only for detecting major cell types. In order to handle this issue, we leverage the post-mortem RNA subgroups by launching OT between the marker matrix *X* and our post-mortem single-cell RNA matrix *Y* ∈ ℝ^*m*×*p*′^. Note that the number *p*^′^of genes in our single-cell RNA data is usually much larger than *p*, so we subset the RNA matrix using only the *p* marker genes from spatial. Once an OT plan has been computed, we use it to transfer the subtypes from post-mortem RNA to spatial data. In order to characterize subgroups that are spatially close in the data, we then create a composition matrix *Z* ∈ ℝ^*n*×*G*^, where *G* is the number of subgroups identified in post-mortem RNA. For each cell *x*, with associated spatial coordinates *c*(*x*) ∈ ℝ^3^, we use the coordinate matrix *C* to identify the cells that are at (spatial) distance at most 60 pixels from *x* :

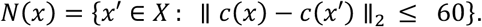

Then, the composition profile of *x* is computed as the fraction of each subgroup in the neighborhood: 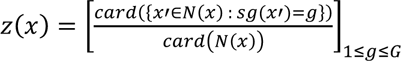 where *sg*(*x*) denote the RNA subgroup of *x* (identified after OT transfer). The composition matrix thus characterizes cells by the composition of their neighborhoods, and can be used for clustering in order to group cells together according to subgroups that are around them. In our analysis, we cluster composition profiles using community detection. More precisely, we first build a *k* -nearest neighbor graph using the Euclidean distances between composition profiles. Then, we run community detection with modularity to partition the nodes into communities. The idea behind modularity is to find a partition of the nodes such that the number of edges induced by the subgraphs formed by the communities is as larger as possible than the expected number of edges of a random graph. More formally, the modularity of a graph *G* = {*V*, *E*} = {(*v*_1_, …, *v*_*n*_), *E*} with a partition of the nodes into *m* communities 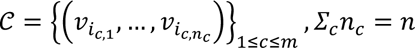, is computed as:

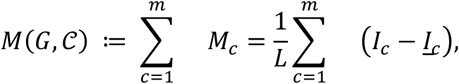

where *I*_*c*_ ≔ *card E*_*c*_, *E*_*c*_ is the set of edges of the subgraph induced by community 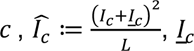 is the set of edges between community *c* and the rest of the nodes, *L* ≔ *I*_*c*_ + 2<u>*I*</u>_*c*_ + *I*<u>_*c*_</u>, *I*<u>_*c*_</u> ≔ *card*(<u>*E*</u>_*c*_), and <u>*E*</u>_*c*_ is the set of edges of the subgraph induced by the nodes outside community *c*. Community detection with modularity amounts to finding a partition 𝒞 that maximizes *M*(*G*, 𝒞). The main advantage of modularity is that it is parameter-free, and thus no tuning is required. For computing such an optimal partition, we use the Louvain algorithm of Blondel et al.^112^, available in the networkx Python package. Finally, we assess the robustness of our partition with respect to the choice of *k* in the construction of the nearest neighbor graph (prior to running community detection). For this, we pick the most stable *k* in the list {5*k*′ : *k*′ ∈ 2, … ,20}, where stability is computed with two indicators:

1. the mean Jaccard similarity 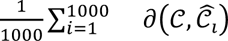 between the current community partition 𝒞 and the community partitions 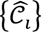 associated to 1000 random subsamples of the data sets, of size 90 % of the total number of cells, and where 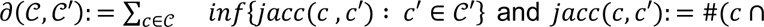 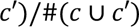.

2. the p-value associated to a two-sample permutation test (computed with 1000 permutations of the composition profile dimensions) on the test statistic measuring the difference between two sets of communities through their Jaccard similarities: 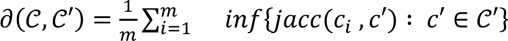 where 𝒞 is a community set with *m* communities.

In order to use these indicators for getting optimal communities, we pick a first estimate of *k* among candidate values with corresponding mean Jaccard similarity above 0.6 and p-value below 0.05 (and we resolve the tie between acceptable candidate values by choosing the value of *k* with the smallest number of communities), and we then merge the associated communities by running hierarchical clustering with Euclidean distance between the communities, that are represented by their mean composition profiles according to the composition matrix *Z*. The dendrogram threshold used for merging the communities is computed using the largest merge distance gap in the dendrogram. This ensures that communities with similar composition profiles are eventually merged into final communities.

#### Cellular Network Interaction Analysis

We performed cell-cell interaction analysis for each community using CellPhoneDB^83^. The expression profiles of conditional cells and their neighboring cells in each community were taking as input. CellPhoneDB was run in the statistical mode, which calculated significance of each interacting pair, reflected by adjusted p-values of permutation tests.

