## Supplemental Methods for "Enhancer Dynamics and Spatial Organization Drive Anatomically Restricted Cellular States in the Human Spinal Cord"

GTCTCGTGGGCTCGGGCTACACAAGATGTGTATAAGAGACAG
ME-B_Calling Card 3
GTCTCGTGGGCTCGGTATCAGCGAGATGTGTATAAGAGACAG
ME-B_Calling Card 4
GTCTCGTGGGCTCGGCTCGCAACAGATGTGTATAAGAGACAG
STAB-N701 Indexing Primer
CAAGCAGAAGACGGCATACGAGATTCGCCTTAGTTCAGACGTGTGCTCGTGGGCTCGG
STAB-N702 Indexing Primer
CAAGCAGAAGACGGCATACGAGATCTAGTACGGTTCAGACGTGTGCTCGTGGGCTCGG
STAB-N703 Indexing Primer
CAAGCAGAAGACGGCATACGAGATTTCTGCCTGTTCAGACGTGTGCTCGTGGGCTCGG
STAB-N704 Indexing Primer
CAAGCAGAAGACGGCATACGAGATGCTCAGGAGTTCAGACGTGTGCTCGTGGGCTCGG
STAB-N705 Indexing Primer
CAAGCAGAAGACGGCATACGAGATAGGAGTCCGTTCAGACGTGTGCTCGTGGGCTCGG
STAB-N706 Indexing Primer
CAAGCAGAAGACGGCATACGAGATCATGCCTAGTTCAGACGTGTGCTCGTGGGCTCGG
STAB-N707 Indexing Primer
CAAGCAGAAGACGGCATACGAGATGTAGAGAGGTTCAGACGTGTGCTCGTGGGCTCGG
STAB-N708 Indexing Primer
CAAGCAGAAGACGGCATACGAGATCCTCTCTGGTTCAGACGTGTGCTCGTGGGCTCGG
STAB-customREAD2 Sequencing Primer
GTTCAGACGTGTGCTCGTGGGCTCGG
STAB-customIndex1 Sequencing Primer
CCGAGCCCACGAGACACACGTCTGAAC

$$J_{R,i}^{(1)} = \max_j Jaccard\left(\tilde{C}_{R,i}, D_{R,j}^{(1)}\right),$$

   $1 \le i \le K_R$. $\tilde{C}_{R,i}$ are the cells in $C_{R,i}$ that appear in the subsampled set.
5. Repeat step 3 and 4 for $N$ times. We set $N$ as 100 in our implementation.
6. For each resolution $R$, calculate the stability score as

$$S_R = \frac{1}{N\sum_{i=1}^{K_R}|C_{R,i}|}\sum_{i=1}^{K_R}\left|C_{R,i}\right|\sum_{n=1}^{N} 1_{\left\{J_{R,i}^{(n)}>0.5\right\}},$$

   which is namely the proportion of maximal Jaccard index greater than 0.5 weighted by cluster size. $\left|C_{R,i}\right|$ is the number of cells in $C_{R,i}$. $1_{\{\cdot\}}$ is the indicator function.

Our observation is that $S_R$ has a general decreasing trend in terms of $R$. We select the $\hat{R}$ corresponding to the elbow point of $S_R$, and the related clusters $C_{\hat{R},i}$, $1 \le i \le \hat{R}$ are taken as the subtypes.

*Cluster Annotation for Donor 1*

Given the obtained subtype labels for cells from freshly resected tissues, we inferred the subtypes for RNA cells from the post-mortem tissues using the optimal transportation plan between RNA-seq data. For each cell in the SCRB RNA-seq data, we went through the values in the optimal transportation plan matrix between that cell and every cell in the 10X Multiome RNA data. We selected the cells in 10X Multiome RNA data that corresponded to the top 20 values, and the most frequent subtype associated with the selected cells was taken as the subtype inference. In a similar way, we inferred subtypes for the ATAC cells from post-mortem tissues using the optimal transportation plan between the SCRB RNA-seq data and Biorad ATAC-seq data.

*Canonical Correlation Analysis*

In this section, we let $X \in \mathbb{R}^{n\times p}$ and $Y \in \mathbb{R}^{m\times p}$ denote the two data sets obtained after data preprocessing [cf Anqi's part], where $n$ and $m$ are the numbers of cells in the first and second data sets respectively, and $p$ is the number of genes. We first embed $X$ and $Y$ in a common, low-dimensional common space $Z$ using Canonical Correlation Analysis (CCA). The objective of CCA is to find a linear projection of the data such that the correlation matrix $C = corr(X, Y)$ is as large as possible, where correlation is defined between two (empirical) random variables $x, y \in \mathbb{R}^p$ as

$$corr(x, y) = \frac{\sum_{i=1}^{p} (x_i - \underline{x})(y_i - \underline{y})}{\sqrt{var(x)var(y)}},$$

and $var(x) = \sum_{i=1}^{p} \left(x_i - \underline{x}\right)^2$. In particular, CCA aims at finding projection matrices $A \in \mathbb{R}^{n\times d}$ and $B \in \mathbb{R}^{m\times d}$, for some low dimension $d \in \mathbb{N}^*$, such that $corr(A^T X, B^T Y)$ is maximized. In practice, we compute the correlation matrix $C \in \mathbb{R}^{n\times m}$ between our two data sets, and we then compute a singular value decomposition of $C$ : $C = A\Sigma B^T$. The left and right singular vectors $A \in \mathbb{R}^{n\times d}$ and $B \in \mathbb{R}^{m\times d}$ of this SVD provide the two embeddings that maximize the correlation. In our analysis, we set $d$ as the dimension for which the explained variance achieves $99$ % of the total variance.

### Optimal Transport

Once CCA has been used to embed $X$ and $Y$ in a common embedding space, we use Optimal Transport (OT) to align the cells. OT is a very common tool of applied mathematics that allows to compare discrete probability measures by finding an alignment, or correspondence, between the support of the measures. More formally, given a space $X$ endowed with a cost function $c: X \times X \to \mathbb{R}_+$, and two discrete measures $\mu$ and $\nu$ on $X$, namely measures that can be written as positive combinations of Dirac measures, $\mu = \Sigma_{i=1}^{n} a_i \delta_{x_i}$ and $\nu = \Sigma_{j=1}^{m} b_j \delta_{y_j}$ with weight vectors $a \in \mathbb{R}_+^n$, $b \in \mathbb{R}_+^m$ satisfying $\Sigma_i a_i = \Sigma_j b_j$ (i.e., the measures have same total masses) and all $x_i, y_j \in X$, the $n \times m$ cost matrix $C = \left(c(x_i, y_j)\right)_{ij}$ and the set of candidate transportation matrices, defined as

$\Pi(a,b) \coloneqq \{P \in \mathbb{R}_+^{n\times m} \ : \ P1_m = a, P^T 1_n = b\}$

define a so-called optimal transport problem. The optimal transport plan $P^*$ can be computed using the following linear program:

$P^* = arg\ arg\ \langle P, C\rangle$ (1)

where $\langle \cdot,\cdot \rangle$ is the Frobenius dot product, i.e., $\langle P, C\rangle = \sum_{i=1}^{n} \sum_{j=1}^{m} P_{ij} C_{ij}$. Unfortunately, it is well-known that solving the optimal transport problem is intractable when data sets are large. In particular, our single–cell data are too large to compute the optimal solution $P^*$ exactly. A common and very efficient workaround[11] is to consider an entropic regularization of the optimal transport problem, namely:

$P_{C,\epsilon}^* = arg\ arg\ \langle P, C\rangle - \epsilon \cdot h(P)$ (2)

where $\epsilon > 0$ and the negentropy $h$ is defined as $h(P) \coloneqq -\Sigma_{ij} P_{ij}\left(log\ log\ (P_{ij} - 1)\right)$. Since the negentropy is strongly convex, the regularized optimal transport problem admits a unique solution, and can be computed efficiently. Indeed, it is known that $P_{C,\epsilon}^*$ takes the following form:

$$P_{C,\epsilon}^* = diag(u^\epsilon) \cdot K \cdot diag(v^\epsilon) \in \mathbb{R}^{n\times m}$$

where $K$ is computed by exponentiating each term of $C$ with $K \coloneqq e^{-\frac{C}{\epsilon}}$, and $(u^\epsilon, v^\epsilon) \in \mathbb{R}^n \times \mathbb{R}^m$ can be computed as the fixed points of the so-called Sinkhorn map: $S: (u, v) \mapsto \left(\frac{a}{K\cdot v}, \frac{b}{K^T \cdot u}\right)$.

Note that these fixed points are the limits of any iterative sequence $(u_{t+1}, v_{t+1}) = S(u_t, v_t)$, which immediately gives an algorithm to estimate $P_{C,\epsilon}^*$, known as Sinkhorn iterations. The Sinkhorn divergence is defined as the transport cost of the optimal regularized plan, $S_C^\epsilon(a,b) \coloneqq \langle P_{C,\epsilon}^*, C\rangle = (u^\epsilon)^T (K \odot C) v^\epsilon$(where $\odot$ denotes the term-wise multiplication), and is known to converge $S_C^\epsilon(a,b) \to \langle P^*, C\rangle$ as $\epsilon \to 0$, and more precisely $P_{C,\epsilon}^*$ converges to the optimal transport plan solution of (1) with maximal entropy. Finally, OT can be generalized to unbalanced OT whenever (2) is augmented with two Kullback-Leibler terms:

$P_{C,\epsilon,\delta}^*(a,b) = arg\ arg\ \langle P, C\rangle - \epsilon \cdot h(P) + \delta \cdot KL(P1_m, a) + \delta \cdot KL(P^T 1_n, b)$ (3)

$$= arg\ arg\ \langle P, C\rangle - \epsilon h(P) + \delta \sum_{i=1}^{n} (P1_m)_i\ log\ log\ \left(\frac{(P1_m)_i}{a_i}\right) + \delta \sum_{j=1}^{m} (P^T 1_n)_j\ log\ log\ \left(\frac{(P^T 1_n)_j}{b_j}\right)$$

where $P$ ranges now over the set $\tilde{\Pi}$ of the positive $n \times m$ matrices. Again, cost (3) can be solved using Sinkhorn iterations. In our analysis, we always use unbalanced OT between our data sets $\tilde{X}$ and $\tilde{Y}$preprocessed with CCA, using the Euclidean pairwise distance matrix $D = \left(\| \tilde{x_i} - \tilde{y_j} \|^2\right)_{ij} \in \mathbb{R}_+^{n\times m}$ between $\tilde{X}$ and $\tilde{Y}$ as the cost matrix $C \coloneqq D$, and uniform weight vectors $a = \left(\frac{1}{n}\right)_i$ and $b = \left(\frac{1}{m}\right)_j$. The entropic and marginal regularizations $\epsilon$ and $\delta$ are chosen in the list

$\{10^q \cdot med(D) \ : \ q \in [-3,-2,-1,0,1,2,3]\}$,

where $med(D)$ is the median of D. More precisely, $\epsilon$ and $\delta$ are chosen as the smallest values in that list such that numerical errors are avoided. OT transportation plans are computed with the POT Python package. Once an optimal transportation plan $P^* \in \mathbb{R}^{n\times m}$ has been computed, we use it to transfer information (such as, e.g., cell types). For a given cell $\tilde{x_i}$, we aggregate the k-th largest values and their indices

$$Ind_k(i) = \left\{ j_{i_1}, \dots, j_{i_k} : P^*_{i,j_{i_q}} \geq P^*_{i,j_{i_{q'}}} \forall q \in \{i_1, \dots, i_k\},\ q' \in \{1, \dots, m\} \setminus \{i_1, \dots, i_k\} \right\}$$

of row $i$ in matrix $P^*$, and select the most frequent information associated to this subset. Moreover, in order to avoid selecting an arbitrary $k$ for transferring information, we run this transfer for $k \in \{5k' : k' \in 2, \dots, 20\}$, and select again the most frequent information among all transferred ones (one for each value of $k$ ).

$$Y = \theta\Phi Y = \theta\Phi$$

where $Y \in R^{S,n}$ is the gene expression matrix with n genes and S samples, $\Phi \in R^{m,k}$ is the TFA with k TFs, and $\Theta \in R^{k,n}$ is the GRN. $\Phi$ is unknown and to deconvolve and find TFA estimates we impose a task specific prior $P_C$ with elements $\in 0, 1$ and use that to solve for an estimate of TFA;

| *Expression* | *H3K27ac coefficient* | *H3K27me3 coefficient* | *H3K4me1 coefficient* | *Class* |
|---|---|---|---|---|
| Down | + | | | Deactivated |
| Down | | - | | Repressed |
| Down | | | - | Deprimed |
| Down | | | + | Primed |
| Up | + | - | * | Activated/Derepressed |
| Up | + | | + | Activated/Primed |
| Up | | - | + | Derepressed/Primed |
| Up | + | | | Activated |
| Up | | - | * | Derepressed |
| Up | | | + | Primed |
| Up | | | - | Deprimed |

In more detail, let

$$P = \begin{pmatrix} Q & R \\ 0 & I \end{pmatrix}$$

be the transition matrix for this absorbing Markov chain. We have closed-form expressions for the various properties of this process (Kemeny and Snell, 1976). For instance, the expected time to absorption (at any absorbing node) is $N1$, where $N = (I - Q)^{-1}$ is the fundamental matrix of the Markov chain. Separating this out into absorption probabilities at each absorbing node, we let

*Community Detection*
In this section, we explain how we get communities of cells in spatial data. We recall that spatial data is given as a cell by marker matrix $X \in \mathbb{R}^{n \times p}$, and a spatial coordinate matrix $C \in \mathbb{R}^{n \times 3}$. Even though cell types can be inferred from marker gene expression in $X$ , the (relatively) small number of markers does not allow for precise assessment of subgroups, and only for detecting major cell types. In order to handle this issue, we leverage the post-mortem RNA subgroups by launching OT between the marker matrix $X$ and our post-mortem single-cell RNA matrix $Y \in \mathbb{R}^{m \times p'}$. Note that the number $p'$of

genes in our single-cell RNA data is usually much larger than $p$, so we subset the RNA matrix using only the $p$ marker genes from spatial. Once an OT plan has been computed, we use it to transfer the subtypes from post-mortem RNA to spatial data. In order to characterize subgroups that are spatially close in the data, we then create a composition matrix $Z \in \mathbb{R}^{n \times G}$, where $G$ is the number of subgroups identified in post-mortem RNA. For each cell $x$, with associated spatial coordinates $c(x) \in \mathbb{R}^3$, we use the coordinate matrix $C$ to identify the cells that are at (spatial) distance at most $60$ pixels from $x$:

$N(x) = \{x' \in X : \| c(x) - c(x') \|_2 \leq 60\}$.

Then, the composition profile of $x$ is computed as the fraction of each subgroup in the neighborhood: $z(x) = \left[\frac{card(\{x' \in N(x) : sg(x') = g\})}{card(N(x))}\right]_{1 \leq g \leq G}$ where $sg(x)$ denote the RNA subgroup of $x$ (identified after OT transfer). The composition matrix thus characterizes cells by the composition of their neighborhoods, and can be used for clustering in order to group cells together according to subgroups that are around them. In our analysis, we cluster composition profiles using community detection. More precisely, we first build a $k$-nearest neighbor graph using the Euclidean distances between composition profiles. Then, we run community detection with modularity to partition the nodes into communities. The idea behind modularity is to find a partition of the nodes such that the number of edges induced by the subgraphs formed by the communities is as larger as possible than the expected number of edges of a random graph. More formally, the modularity of a graph $G = \{V, E\} = \{(v_1, \dots, v_n), E\}$ with a partition of the nodes into $m$ communities $\mathcal{C} = \left\{\left(v_{i_{c,1}}, \dots, v_{i_{c,n_c}}\right)\right\}_{1 \leq c \leq m}$, $\Sigma_c n_c = n$, is computed as:

$$M(G, \mathcal{C}) := \sum_{c=1}^{m} M_c = \frac{1}{L} \sum_{c=1}^{m} \left(I_c - \underline{I_c}\right),$$

where $I_c := card(E_c)$, $E_c$ is the set of edges of the subgraph induced by community $c$, $\widehat{I_c} := \frac{(I_c + \underline{I_c})^2}{L}$, $\underline{I_c}$ is the set of edges between community $c$ and the rest of the nodes, $L := I_c + 2\underline{I_c} + I_{\underline{c}}$, $I_{\underline{c}} := card(\underline{E_c})$, and $\underline{E_c}$ is the set of edges of the subgraph induced by the nodes outside community $c$. Community detection with modularity amounts to finding a partition $\mathcal{C}$ that maximizes $M(G, \mathcal{C})$. The main advantage of modularity is that it is parameter-free, and thus no tuning is required. For computing such an optimal partition, we use the Louvain algorithm of Blondel et al.[20], available in the networkx Python package. Finally, we assess the robustness of our partition with respect to the choice of $k$ in the construction of the nearest neighbor graph (prior to running community detection). For this, we pick the most stable $k$ in the list $\{5k' : k' \in 2, \dots, 20\}$, where stability is computed with two indicators:
1. the mean Jaccard similarity $\frac{1}{1000} \sum_{i=1}^{1000} \partial\left(\mathcal{C}, \widehat{\mathcal{C}}_i\right)$ between the current community partition $\mathcal{C}$ and the community partitions $\{\widehat{\mathcal{C}}_i\}$ associated to $1000$ random subsamples of the data sets, of size $90$ % of the total number of cells, and where $\partial(\mathcal{C}, \mathcal{C}') := \sum_{c \in \mathcal{C}} inf\{jacc(c, c') : c' \in \mathcal{C}'\}$ and $jacc(c, c') := \#(c \cap c')/\#(c \cup c')$.
2. the p-value associated to a two-sample permutation test (computed with $1000$ permutations of the composition profile dimensions) on the test statistic measuring the difference between two sets of communities through their Jaccard similarities: $\partial(\mathcal{C}, \mathcal{C}') = \frac{1}{m} \sum_{i=1}^{m} inf\{jacc(c_i, c') : c' \in \mathcal{C}'\}$, where $\mathcal{C}$ is a community set with $m$ communities.

In order to use these indicators for getting optimal communities, we pick a first estimate of $k$ among candidate values with corresponding mean Jaccard similarity above 0.6 and p-value below 0.05 (and we resolve the tie between acceptable candidate values by choosing the value of $k$ with the smallest number of communities), and we then merge the associated communities by running hierarchical clustering with Euclidean distance between the communities, that are represented by their mean composition profiles according to the composition matrix $Z$. The dendrogram threshold used for merging the communities is computed using the largest merge distance gap in the dendrogram. This ensures that communities with similar composition profiles are eventually merged into final communities.

*Cellular Network Interaction Analysis*

We performed cell-cell interaction analysis for each community using CellPhoneDB[21]. The expression profiles of conditional cells and their neighboring cells in each community were taking as input. CellPhoneDB was run in the statistical mode, which calculated significance of each interacting pair, reflected by adjusted p-values of permutation tests.
